# Type 1 immunity linked to CMV exposure improves long-term survival in sepsis without shock

**DOI:** 10.64898/2026.10.13.758958

**Authors:** Wei Qi, Nuzat Zaman, Alexis Rooney, Caroline B.M. Porter, Konstantinos Boulias, Sarah Naz-McLean, Joshua Pickard, Marla Glean, Ndongwa Pemba, Kastin Pan, Shilei Li, Hannah H. Laibinis, Con Moran, Erin Connolly, Grace Valentine, Ramnik J. Xavier, Deborah T. Hung, Lisa Cosimi, Ann E. Woolley, Jacques Deguine

## Abstract

Dysregulated immune responses are at the core of sepsis pathology and can drive both acute organ dysfunction and long-term morbidity and mortality. To better understand the association between specific immune histories, perturbations and sepsis outcomes, we combined immune phenotyping, functional cytokine production assays and antibody repertoire profiling. While sepsis immune profiles were highly heterogeneous, we identified a strong association between the frequency of CXCR3-expressing CD4 memory T cells at time of enrollment and six-month survival in sepsis without shock. Surprisingly, the same population was linked to improved organ function trajectories in septic shock patients but did not influence six-month survival, highlighting the importance of considering physiological states to define immune predictors. Finally, phage display profiling revealed that these Th1 cells were generally expanded in CMV-seropositive individuals, suggesting that previous and latent infections may modulate sepsis outcomes.

## Introduction

Sepsis and septic shock are characterized as a syndrome of organ dysfunction driven by a dysregulated host immune response ^1–3^. Despite advances in clinical care, the mortality and morbidity associated with sepsis remain high, both during the acute phase of the disease and in the following year. Current standard of care primarily focuses on early diagnosis and rapid initiation of appropriate antimicrobial and supportive hemodynamic and pulmonary care ^4^.

While the essential role of immune responses in sepsis has been recognized, the immunobiology of sepsis is complex and highly heterogeneous. Disease in animal models appears primarily driven by inflammatory cytokines such as TNF and IL-1, but human trials of blocking agents have shown no consistent benefit ^5–7^. Interestingly, a recent trial that stratified patients based on hyperinflammation (ferritin levels) or immune paralysis (MHC class II expression on dendritic cells (DCs)) showed that appropriate immune modulation (IL-1 blockade or recombinant IFN-γ, respectively) improved organ dysfunction scores ^8^. While treatment did not impact 28-or 90-day mortality, this study provides a rationale for using immune profiles of septic patients to define proper immune modulation strategies ^9^.

The recognition of this heterogeneity has motivated multiple studies aimed at defining sepsis endotypes, particularly through the use of bulk transcriptomics. These studies led to the identification of factors associated with outcomes, where a more adaptive signature is generally favorable and an innate or coagulopathic signature is associated with increased mortality ^10–13^. Some of these signatures also appear to predict response to treatment, with specific endotypes showing higher mortality upon corticosteroid treatment ^11^. Most studies focused on endotypes at the time of enrollment, which is generally around peak disease severity, but a recent longitudinal study showed that individual trajectories are also variable and may confound endotype classification ^14^. While whole blood gene expression-based profiling offers an integrated view of PBMC composition and states, this method can conflate changes in cell frequencies, for example due to lymphopenia, and gene expression. Additionally, gene expression can be an incomplete proxy of immune cell function and functional profiling of response to stimuli has demonstrated a greater than expected variability across even healthy subjects, a phenomenon that is driven in part by previous exposures like cytomegalovirus infection ^15–17^. Lastly, it remains unknown how systemic physiologic changes in the septic host, such as the development of shock, impact immune cell function and subsequent endotype classification.

Here, we aimed to define the immune states of a cohort of sepsis-at-risk individuals through complementary assays including flow cytometry, immune stimulation and quantification of cytokine production, and antibody repertoire analysis as a proxy of previous exposures. As expected, immune phenotyping showed an enrichment of myeloid cells and relative depletion of lymphocytes, but a high degree of heterogeneity across sepsis patients. While the frequency of main lineages were not associated with disease features or outcomes, we found that the frequency of CXCR3-expressing memory CD4 T cells was positively correlated with 6-month survival in septic individuals without shock, but that this correlation was lost in septic shock.

Stimulation assays similarly showed an association between IFN-γ production and 6-month survival in sepsis without shock, supporting a protective effect of type 1 responses in these patients. Conversely, type 1 responses were associated with increased likelihood of organ dysfunction score improvement in the context of shock but no 6-month survival benefit. Finally, antibody profiling demonstrated an association between anti-CMV antibodies and Th1 abundance during peak illness, suggesting that CMV pre-exposure is a potential driver of immune variability at sepsis presentation. Altogether, this framework suggests that functional immune assays can delineate subsets of patients with distinct long-term outcomes and highlight type 1 responses as a potential lever for intervention in these groups.

## Results

### Immune profiling of patients with suspected sepsis

To characterize the peripheral blood immune landscape of patients at risk for sepsis, we prospectively enrolled hospitalized adults with suspected sepsis (n = 89). Patients were identified by evidence of systemic inflammation (≥2 Systemic Inflammatory Response Syndrome [SIRS] criteria) concurrent with recent microbiologic sampling and/or initiation of antimicrobial therapy, as ascertained by electronic medical record (EMR) review (see **Methods**). Peripheral blood was collected at enrollment (day 0) and at day 7 for PBMC cryopreservation and serum isolation. Using day 14 data, each patient was adjudicated against Sepsis-3 research criteria for the presence of infection and organ dysfunction. Patients were assigned to one of four categories: infected septic (n = 65; comprising patients with septic shock, n = 29, and those without shock, n = 36), infected but not septic (n = 8), non-infectious SIRS (n = 13), and uninfected/possible SIRS (n = 3) (**Figure 1A**). Across three experimental batches, PBMCs were thawed and split to allow for phenotyping by spectral flow cytometry and parallel functional characterization by *ex vivo* stimulation and cytokine quantification.

**Figure 1.**
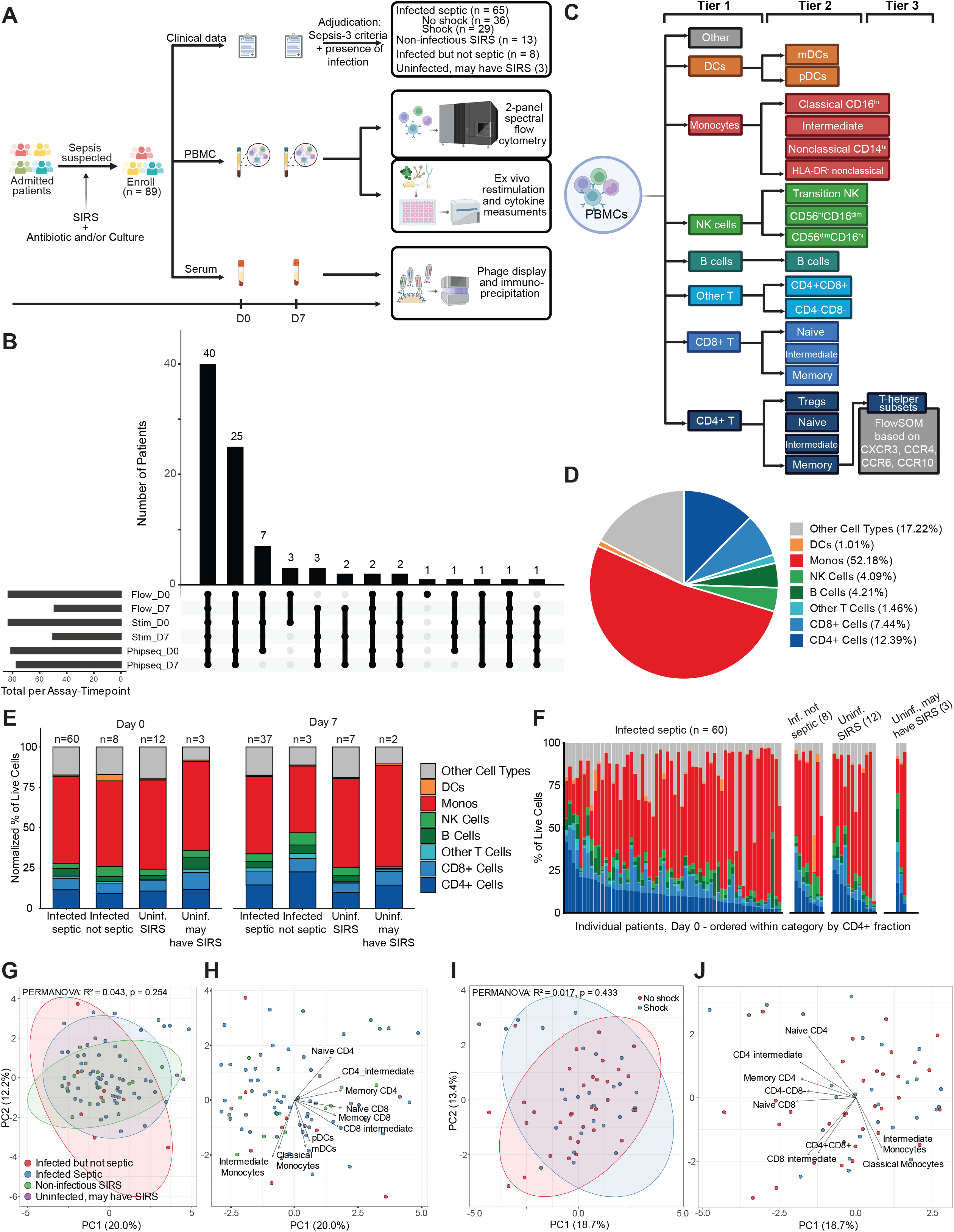
Study design and compositional analysis of PBMCs across adjudicated categories. (A) Prospective cohort design and immune profiling workflow. (B) UpSet plot of assay coverage by timepoint across the 89 cohort patients. Set sizes are Flow D0: 83, Flow D7: 49, Stim D0: 83, Stim D7: 50, PhIP-Seq D0: 81, PhIP-Seq D7: 77; the largest intersection is the 40 patients with all six assay-timepoint combinations. (C) Schematic for gating hierarchy. Tier 1 comprises the eight major lineages used for the composition analyses; Tier 2, 19 sublineages; Tier 3, FlowSOM metaclusters of the memory-CD4 compartment resolved by chemokine receptors (CXCR3, CCR4, CCR6, CCR10). (D) Average Tier 1 lineage composition across all flow samples, expressed as a percentage of live PBMCs. (E) Average Tier 1 lineage composition by adjudicated category at day 0 and day 7 (unpaired), normalized to 100% of live PBMCs, with the number of samples contributing to each bar. (F) Per-patient Tier 1 lineage composition at day 0. One column per patient, faceted by adjudicated category and ordered within category by CD4+ fraction. (G) Principal component analysis (PCA) of day-0 Tier 1 lineage composition (as % live PBMCs) across all four adjudicated categories (sepsis-at-risk with day-0 samples, n = 83). PC1 and PC2 are shown with 95% confidence ellipses. Adjudicated categories explain no detectable share of the variance (PERMANOVA R² = 0.043, p = 0.254). (H) PCA of day-0 Tier 1 lineage composition with loading vectors for the five strongest contributors to each axis. (I) PCA of day-0 Tier 1 lineage composition of infected septic patients (n = 60), colored by septic shock status. PC1 and PC2 are shown with 95% confidence ellipses. Shock status does not explain Tier 1 compositional variance (PERMANOVA R² = 0.017, p = 0.433). PCA of day-0 Tier 1 lineage composition of infected septic patients with loading vectors for the five strongest contributors to each axis.

Separately, serum samples were profiled by phage-display and immunoprecipitation sequencing (PhIP-Seq) to map individual antibody repertoires. Most patients had all longitudinal (n = 40) or day 0 (n = 37) samples profiled by all three modalities, while a small number of patients (n = 12) were profiled by only a subset of assays (**Figure 1B**).

We first assessed PBMC composition in sepsis-at-risk patients. Using separate T cell and B cell/myeloid panels, cells were resolved into parent lineages (Tier 1, 8 populations) and their respective subsets (Tier 2, 19 populations) (**Figure 1C, Figure S1**). At day 0, live PBMCs were dominated by monocytes across all four categories (infected septic, median 56.0%, [IQR 41.3-66.8]; infected but not septic, 56.9%, [IQR 48.1-67.6]; non-infectious SIRS, 62.3%, [IQR 38.6-73.3]; uninfected/possible SIRS, 60.4%, [IQR 40.3-72.4]; **Table S1**), reflecting the fact that all samples were collected in the context of critical care. CD4+ T cells were the most abundant lymphocyte population, comprising 9.9% (IQR 6.0-13.7), 7.8% (IQR 4.8-13.3), 8.3% (IQR 5.7-16.1), and 11.6% (IQR 8.5-14.7) of live PBMCs in infected septic, infected but not septic, non-infectious SIRS, and uninfected/possible SIRS patients, respectively (**Figure 1D, Table S1**). At day 0 and day 7, Tier 1 lineage composition did not differ significantly between adjudicated categories, although within-category composition exhibited significant heterogeneity (**Figure 1E-F, Table S1**). Paired day 0 and day 7 samples were available for infected septic (n = 32) and non-infectious SIRS (n = 6) patients, in whom Tier 1 composition was broadly stable over the first week; paired samples from infected but not septic (n = 3) and uninfected (n = 2) patients were not tested formally given the low sample number (**Figure S2A**). We do however note that attrition prior to day 7 was primarily due to early discharge, potentially representing a less severe subset of patients. Given the observed Tier 1 compositional stability between day 0 and day 7, and the incomplete day 7 sampling, we prioritized analyses of Day 0 samples.

Importantly, we first aimed to establish whether Day 0 - i.e. day of enrollment - was generally representative of peak of disease by examining the timing of maximal organ dysfunction, a defining feature of sepsis and a major contributor to its mortality. Among infected septic patients (n = 65), the day of peak Sequential Organ Failure Assessment (SOFA) score was day 0 in the median patient (IQR-1 to 0 days; **Figure S2B**). Day 0 data therefore captured patients at or immediately after the height of organ dysfunction, the point at which a prognostic biomarker would be valuable for clinical decision-making.

We next asked whether finer-grained composition at enrollment tracked with adjudicated clinical categories. Because Tier 2 spanned 19 sublineages, we performed a principal component analysis (PCA) of day 0 frequencies from all sepsis-at-risk patients (n = 83 with day-0 compositional data). PC1 and PC2 captured 20.0% and 12.2% of the variance respectively, and PC3-10 each captured less than 10% of total variance (**Figure S2C**). PC1 was dominated by the T cell compartment, with T cell lineages loading opposite monocyte subsets, while PC2 separated classical and intermediate monocytes and DC subsets from NK cells, HLA-DR-nonclassical monocytes, and naive T cells (**Figure 1H, Figure S2D**). Adjudicated category explained 4.3% of the variance in Tier 2 composition and did not define separable clusters (PERMANOVA R² = 0.043, p = 0.254, **Figure 1G**). Focusing specifically on the infected sepsis subgroup at day 0 (n = 60), we again observed broad heterogeneity in compositional profiles; shock status did not separate patients by Tier 2 composition (PERMANOVA R² = 0.017, p = 0.433; **Figure 1I**). PC1 and PC2 (18.7% and 13.4% of variance) similarly described a myeloid-lymphoid gradient (**Figure 1J**, **Figure S2E**). We also examined four additional variables in the infected septic group at day 0, including age, sex, BMI, and anatomic source of infection, none of which was associated with Tier 2 composition (**Figure S2F**). Thus, the dominant axes of variation in the circulating immune compartment were shared across all patients with suspected sepsis and were not organized by clinical categories or by the presence of shock. Moreover, both Tier 1 and 2 immune compositions demonstrated a high degree of within-category heterogeneity, suggesting that lineage-level composition may not be sufficient to inform clinical outcomes.

### CXCR3+ memory CD4+ T cell abundance at enrollment associates with 180-day survival in sepsis without shock

Shock status and survival are directly related, as shock is associated with higher in-hospital mortality. Indeed, in our cohort, 180-day survival was 72% (95% CI, 55-84%) in those without shock (n = 36) and 55% (95% CI, 36-71%) in those with shock (n = 29) (log-rank p = 0.12; **Figure 2A, Table S2**). The separation began in the acute phase, where mortality by day 14 was 14% (4/29) in those with shock, and only 6% (2/36) in those without shock; by day 30 mortality was 24% (7/29) and 11% (4/36), respectively. The direction and magnitude of the survival difference are consistent with the established mortality of sepsis with and without shock, and we therefore stratified subsequent analyses by shock status. Notably, with no loss of follow-up before 180 days, survival in both groups continued to decline beyond the first month (6 of 10 deaths without shock and 6 of 13 with shock occurred after day 30), indicating that mortality accrues well beyond the acute phase.

**Figure 2.**
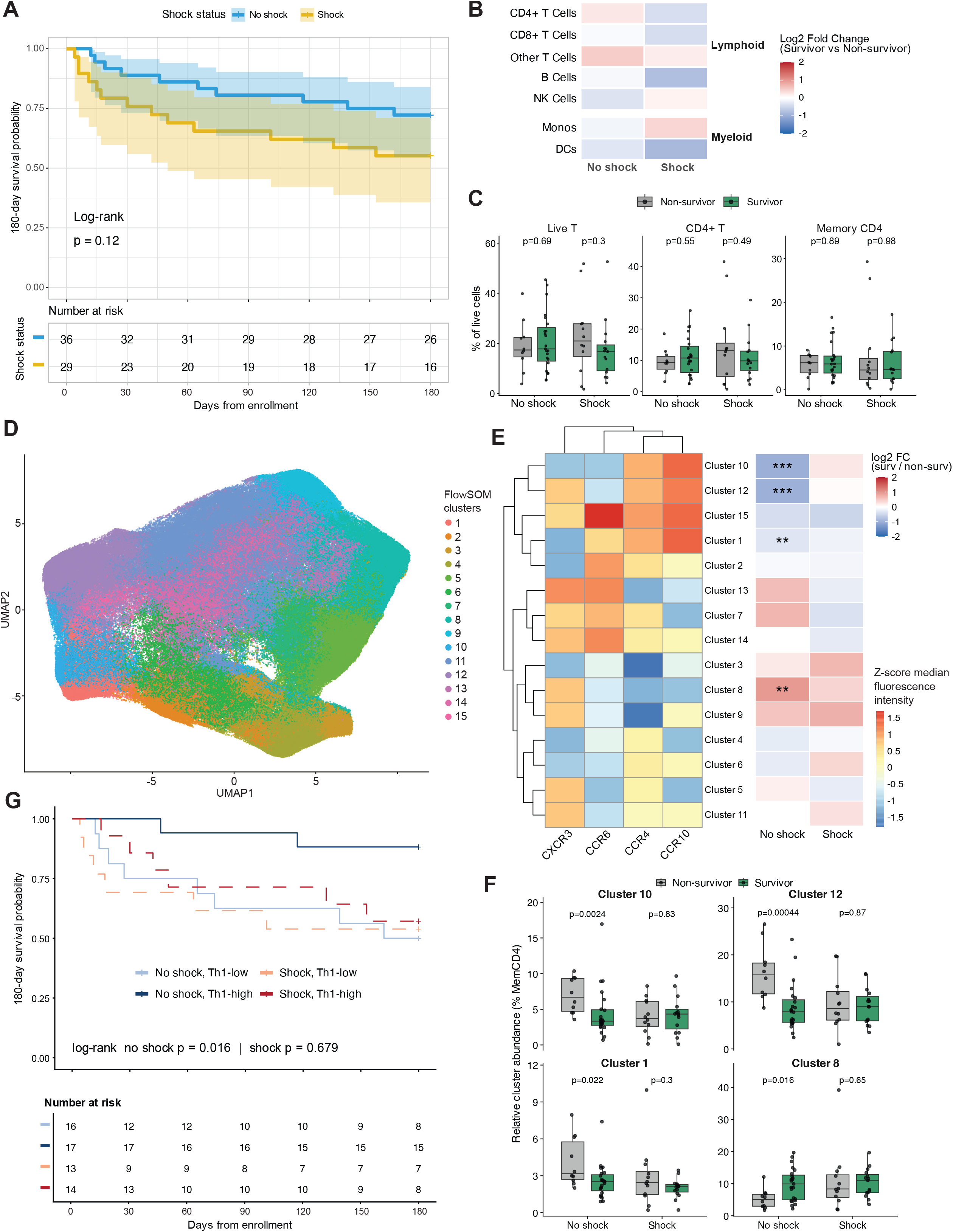
Memory CD4+ T cell helper subset analysis defines populations associated with 180-day survival in sepsis without shock. (A) Kaplan-Meier estimates of 180-day survival among infected septic patients by septic shock status (no shock n = 36, shock n = 29), with log-log confidence bands and the number at risk indicated in table below. Statistical significance was determined by the log-rank test, p value as shown. (B) Tier 1 lineage log2 fold change in day-0 abundance, survivors versus non-survivors, stratified by shock (no shock survivors n = 23, no shock non-survivors n = 10; shock survivors n = 15, shock non-survivors n = 12). Wilcoxon rank-sum with Benjamini-Hochberg correction for multiple comparisons across the 7 lineages per stratum. Significance defined as *** FDR < 0.05, ** FDR < 0.1, * FDR < 0.25. (C) Boxplots of the nested hierarchy from T-cell to memory CD4+ T-cell compartments, as a percentage of live PBMCs, by survival within shock stratum. Wilcoxon rank-sum, unadjusted p values indicated. (D) UMAP embedding of memory CD4+ T-cells colored by FlowSOM clusters (k = 15). Unsupervised FlowSOM clustering performed on CXCR3, CCR4, CCR6 and CCR10 fluorescence intensities. (E) Left: z-scored median fluorescence intensity of the 4 chemokine receptors for each FlowSOM cluster, rows and columns ordered by hierarchical clustering. Right: log2 fold change in day-0 cluster abundance (% of memory CD4) between survivors and non-survivors within each shock stratum. Wilcoxon rank-sum with Benjamini-Hochberg correction for multiple comparisons across the 15 clusters per stratum. Significance defined as *** FDR < 0.05, ** FDR < 0.1, * FDR < 0.25. (F) Boxplots of day-0 abundance of each FlowSOM cluster reaching FDR < 0.25 in (E): Clusters 1, 8, 10 and 12, as a percentage of memory CD4, by survival within shock stratum (Wilcoxon rank-sum, unadjusted p shown). (G) Kaplan-Meier estimates of 180-day survival by day-0 Cluster 8 (CXCR3+ canonical Th1 cluster) abundance (% of memory CD4) in infected septic patients, split at the median within each shock stratum into Th1-high (dark blue/dark red) and Th1-low (light blue/light orange). Sepsis without shock (solid lines), septic shock (dotted lines). Table below indicates the number of patients at risk at 30-day intervals. Statistical significance was determined by the log-rank test, p values as shown.

Hypothesizing that physiologic derangements in septic shock may alter the immune drivers of survival, we compared the day 0 immune parameters between sepsis survivors and non-survivors, stratified by shock status. No lineage differed significantly in either stratum (**Figure 2B**). Among the 19 Tier 2 sublineages, only CD56^bright^CD16^dim^ NK cells showed a differential trend in the shock stratum (median 0.04% vs 0.12% of live PBMCs in survivors vs non-survivors; FDR = 0.11; **Figure S3A**); and overall Tier 2 composition did not separate survivors from non-survivors (no shock: PERMANOVA R² = 0.029, p = 0.498; shock: R^2^ = 0.031, p = 0.684; **Figure S3B**), this high heterogeneity was for example reflected across T cell population, either total, CD4^+^ or CD4^+^ memory (**Figure 2C**). Given the functional diversity of memory CD4^+^ T cells, we next employed unsupervised FlowSOM clustering based on surface chemokine receptor fluorescence intensities (Tier 3: CXCR3, CCR4, CCR6, and CCR10), using these chemokine receptors as surrogates of functional state ^18,19^ to define 15 clusters with varying combinations of chemokine receptor expression (**Figure 2D**).

Strikingly, in sepsis without shock, a Th1-polarized cluster (CXCR3+CCR4-, Cluster 8) was significantly enriched in survivors, whereas Th2-polarized clusters (CCR4+CCR10+, Clusters 1, 10, and 12) were enriched in non-survivors (FDR < 0.1 across 15 clusters; **Figure 2E-F**); conversely, no cluster differed significantly between survivors and non-survivors of septic shock. To further evaluate the association between Th1-polarized cluster abundance and survival, we constructed Kaplan-Meier survival curves comparing patients with high versus low Th1 cluster abundance, defined by the median abundance within each shock stratum. Among patients without shock (n = 33 with Day 0 cytometry data), 180-day survival diverged based on Th1 cluster abundance at enrollment. Patients with above-median Th1 abundance demonstrated significantly higher 180-day survival than those below the median (88% [95% CI, 61-97%] vs 50% [95% CI, 25-71%]; log-rank p = 0.016; **Figure 2G**; **Table S2b**). In a univariate Cox model, above-median Th1 cluster abundance was associated with an 82% reduction in the hazard of death (HR 0.18, 95% CI 0.04-0.87, p = 0.032). By contrast, Th1 abundance had no significant impact on mortality in individuals with shock (HR 0.79, 95% CI 0.25-2.45, p = 0.680). While the total number of deaths (n = 10 in no shock group) limited multivariable modeling, separate sensitivity models confirmed that the Th1 hazard ratio remained essentially unchanged after adjustment for age, maximum SOFA, or immunocompromised status (Table S2b). Altogether, these analyses show that in our cohort, a higher day-0 abundance of CXCR3+ memory CD4+ T cells (Cluster 8) at time of enrollment is associated with improved 6-month survival in sepsis without shock, providing an early predictor of post-acute mortality risk.

### CXCR3+ memory CD4+ T cell abundance tracks organ dysfunction recovery in septic shock

To understand the relationship between immune cell composition and organ dysfunction in sepsis, we turned to Sequential Organ Failure Assessment (SOFA) scores. Daily maximum SOFA scores were computed using values collected from the Electronic Health Record (EHR), missing component scores were imputed as described (see **Methods**). To evaluate SOFA trajectory, we calculated the mean SOFA score across days 1 to 8, and defined “improvers” and “non-improvers” by an absolute threshold of whether that mean fell by at least 1.4 points from baseline (day 0). The eight-day assessment window and 1.4-point threshold were selected based on the ImmunoSep trial design ^8^, which was informed by prior clinical trials ^20,21^ demonstrating an association between post-enrollment day 8 SOFA and 28-day mortality.

Subjects stop contributing to the trajectory once their index admission ends by discharge alive, death or transition to CMO; as such, group medians at later days are computed on progressively fewer patients, as shown in the histogram (**Figure 3A**).

**Figure 3.**
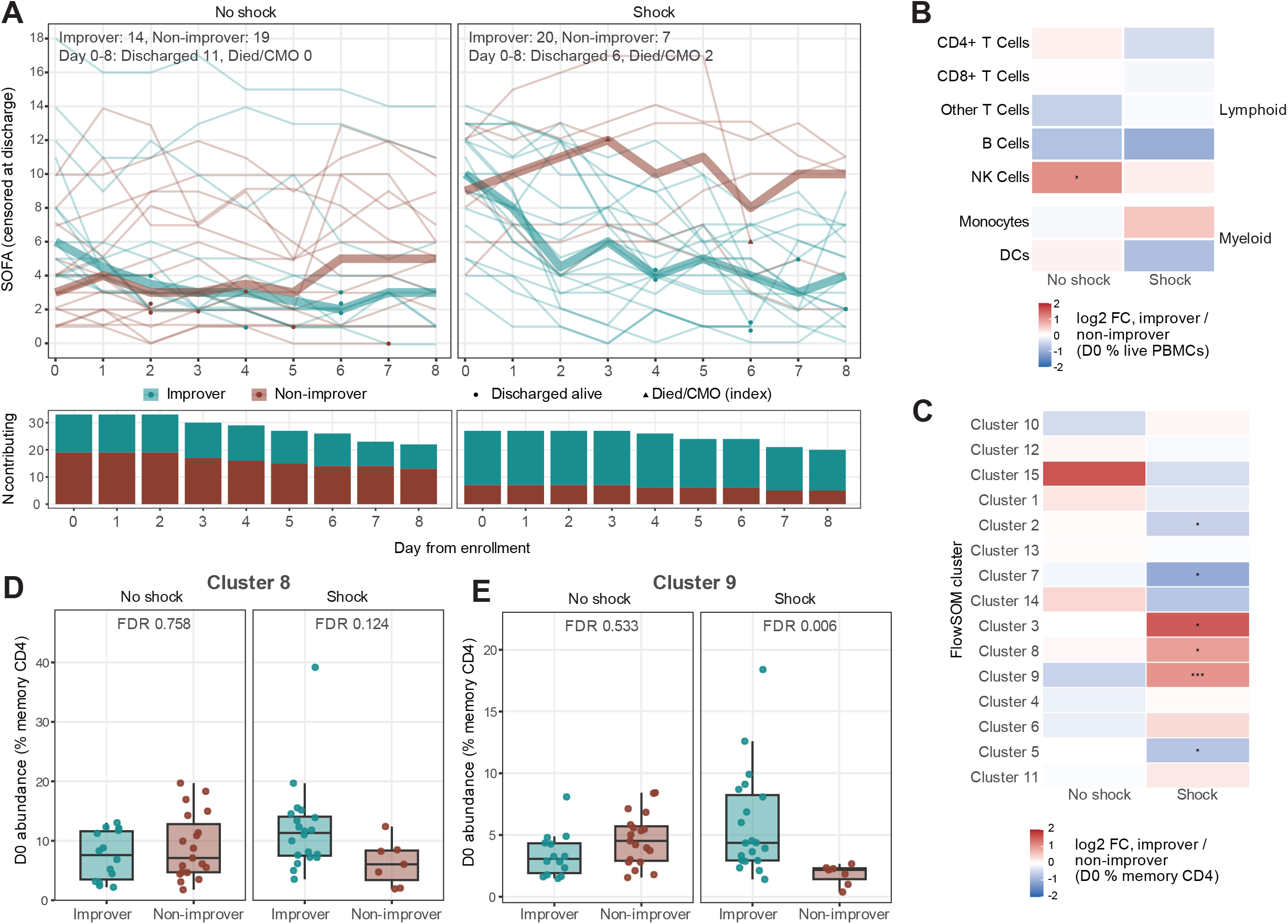
Day 0 CXCR3+ memory CD4+ T cell abundance is associated with organ function trajectory in septic shock. (A) Daily SOFA trajectories over days 0-8 for each infected septic patient with day-0 flow sample, faceted by shock status and colored by improver status under the absolute definition, with the per-group median (thick lines) and the number of patients still contributing each day shown below. Without shock improvers n = 14, non-improvers n = 19; with shock improvers n = 20, non-improvers n = 7. Trajectories are censored at discharge as alive (circle) or as deceased/transitioned to CMO (triangle). (B) Tier 1 lineage log2 fold change in day-0 abundance (% of live PBMCs), SOFA improvers versus non-improvers defined by absolute definition, within shock stratum. Benjamini-Hochberg correction for multiple comparisons across the 7 lineages per stratum. Significance defined as *** FDR < 0.05, ** FDR < 0.1, * FDR < 0.25. (C) FlowSOM cluster log2 fold change in day-0 abundance (% of memory CD4), SOFA improvers versus non-improvers defined by absolute definition. Benjamini-Hochberg correction for multiple comparisons across the 15 clusters per stratum. Significance defined as *** FDR < 0.05, ** FDR < 0.1, * FDR < 0.25. (D) (E) Boxplots of day-0 abundance of (D) Cluster 8 and (E) Cluster 9 as a percentage of memory CD4, by absolute improver status within shock stratum. BH-FDR as shown.

Among 60 septic patients with a day-0 sample (33 without shock, 27 with shock), two patients died or transitioned to CMO between enrollment and day 8, both in the shock stratum; 17 patients were discharged alive within that window (n = 11 without shock, n = 6 with shock).

Applying the absolute definition, 14 of 33 patients without shock were classified as improvers, with median SOFA of 6 (IQR 4.25-10.25) at enrollment falling to 3 (IQR 1-6) by day 8 (n = 9 still contributing); while 19 of 33 were classified as non-improvers, this subgroup showed a lower SOFA of 3 (IQR 2-6) at time of enrollment, rising to 5 (IQR 3-7) by day 8 (**Figure 3A, Table S3**). In the shock group, improvers (n = 20) declined from 10 (IQR 7.75-12) at baseline to 4 (IQR 2-6.5) by day 8 (n = 15), whereas non-improvers (n = 7) showed no decline, at 9 (IQR 7-12) and 10 (IQR 4-11), respectively. Given the differential trajectories of organ dysfunction scores, we next asked whether subset abundance was associated with organ dysfunction.

We began by assessing whether abundance at Tier 1 parent and Tier 2 sublineage levels tracked with baseline organ dysfunction. No population showed more than a weak association with baseline SOFA; the strongest was a negative correlation for B cells in patients without shock (rho =-0.42, FDR 0.102; **Figure S4A-B**). We then turned to the change in organ dysfunction over time, focusing on SOFA improvement as defined by the absolute definition. At the Tier 1 parent level, there was no significant difference between major lineages except NK cells, which were found to be only nominally more abundant in non-shock SOFA improvers under the absolute definition (FDR = 0.20, **Figure 3B**, **Table S3b**); at the sublineage level, there was no significant difference between subset abundances of improvers and non-improvers (**Figure S4C**). However, comparing FlowSOM cluster abundance within the memory CD4+ compartment, 6 of 15 clusters differed between improvers and non-improvers in the shock stratum under the absolute definition (FDR < 0.25); three were more abundant in improvers (Clusters 3, 8 and 9) and three less abundant (Clusters 2, 5 and 7; **Figure 3B**). Clusters 8 and 9 are both CXCR3-expressing Th1 clusters (**Figure 2E**), differing primarily by CCR10 expression, a marker associated with skin and mucosal tissue homing. Both clusters were more abundant in improvers within the shock stratum (Cluster 8: absolute FDR = 0.124; Cluster 9: absolute FDR = 0.006; **Figure 3D-E**; **Table S3c**).

Under the same absolute definition, we did not observe any significant changes in patients without shock, although we note that because of lower baseline SOFA scores, a floor effect limited the ability of subjects to achieve a 1.4-point decrease. Indeed, we did not observe a meaningful separation in SOFA trajectories between no-shock improvers and non-improvers (**Figure 3A**), potentially explaining the lack of immunological differences between these subgroups. Altogether, this integration of organ dysfunction trajectories and immune parameters suggest that higher abundance of CXCR3-expressing memory CD4+ T cells is associated with organ recovery in septic shock. This finding complements our observation of increased 6-month survival associated with the same populations is sepsis without shock, and suggests that these cells are generally associated with improved disease parameters, although this manifests differently in the very distinct physiological contexts or sepsis without or with shock.

### T-cell IFNɣ capacity scales with memory CD4 abundance in sepsis without shock

To build upon the phenotypic characterizations of circulating PBMCs, we next assessed the functional capacity of PBMCs from sepsis-at-risk patients through *ex vivo* stimulation. We stimulated the day 0 and day 7 samples from infected septic (day 0, n = 59; 1 removed from the cohort due to insufficient cell count; day 7, n = 35) and infected but not septic patients (day 0, n = 8) with a panel of seven stimuli, including Toll-like receptor (TLR) ligands (LPS, R848), a pan-T cell activator (anti-CD3/CD28), a cytokine (IL-1β) and heat-killed whole microbes (HKSA, HKEB, HKCA). After 24 hours, supernatants were collected and we quantified the production of 12 distinct cytokines. We also included PBMCs from independently collected healthy donors (n = 4) in this analysis, although we note that these were collected following different procedures and serve as a technical reference rather than a matched biological comparator. Innate responses to microbial stimulation, such as TNF-α induction by LPS, R848, IL-1β, or heat-killed *E. coli,* were similar across healthy donors, infected non-septic, and infected septic subjects (**Figure 4A, Figure S5A**), confirming that PBMCs from infected patients were in a broad sense normally responsive *ex vivo*. However multiple stimuli or output cytokines showed severe defects in sepsis.

**Figure 4.**
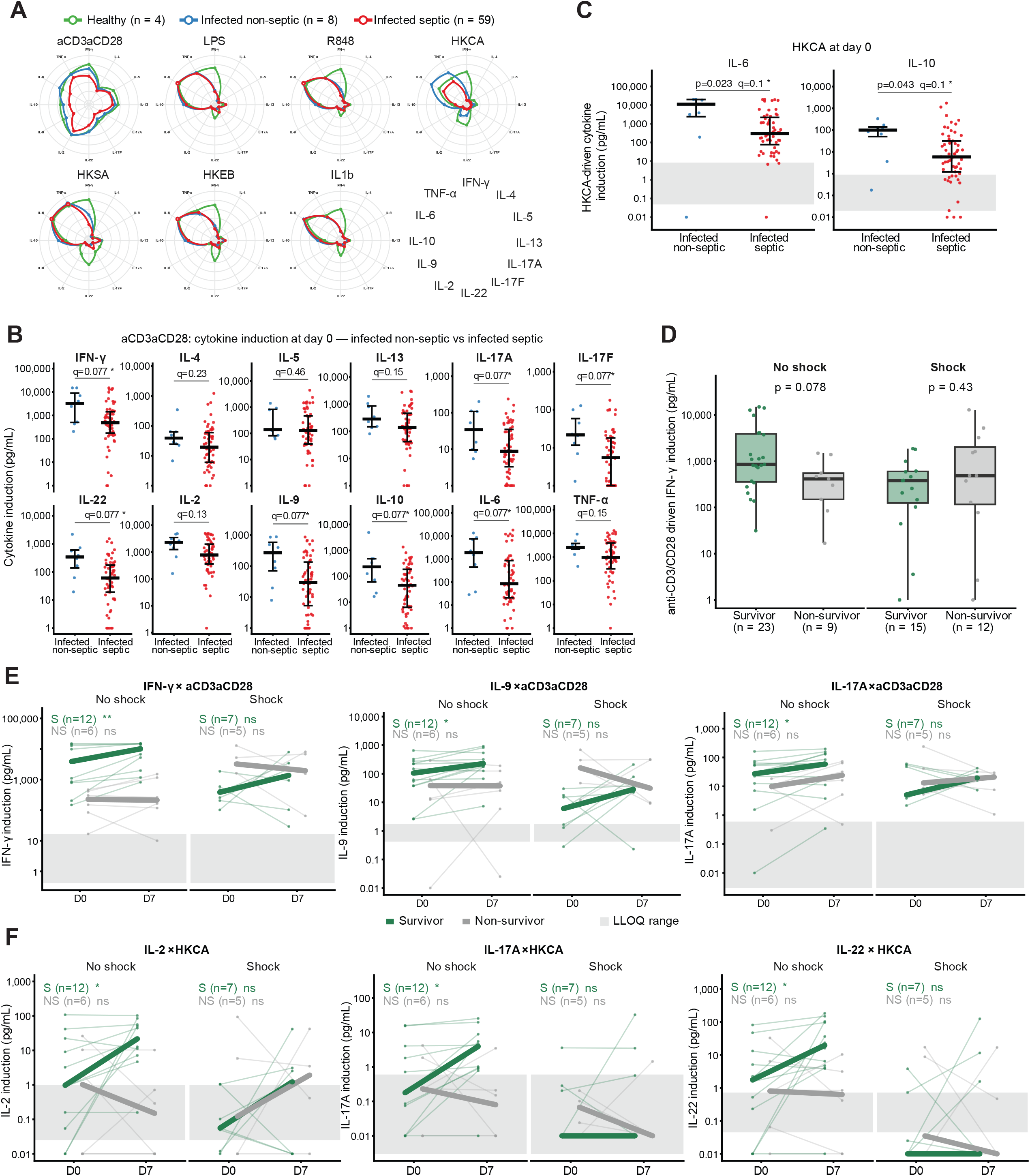
Analysis of PBMC responsiveness to stimulation in infected septic subjects. (A) Radar plot summary of the 12-cytokine induction profile for each of 7 stimuli (anti-CD3/CD28, LPS, R848, heat-killed C. albicans, S. aureus, E. coli, and IL-1β), colored by healthy donors, infected non-septic, and infected septic (day 0 samples from the latter two groups). Open circles indicate responses with >50% of samples at or above the upper limit of detection. Cytokine legend is indicated in the bottom right corner, clockwise from top: IFNɣ, IL-4, IL-5, IL-13, IL-17A, IL-17F, IL-22, IL-2, IL-9, IL-10, IL-6, TNF-ɑ. Each ring represents 1 log(10) change in induction, from a value of 100,000 (outer ring) to 1 (middle). (B) Dotplots of day-0 responses to anti-CD3/CD28, separated by infected septic and infected-nonseptic patients. Median and interquartile range per group as indicated. (C) Dotplots of day-0 innate responses to heat-killed *C. albicans*, separated by infected septic and infected-nonseptic patients. Median and interquartile range per group as indicated. (D) Boxplots of day-0 IFNγ induction by anti-CD3/CD28 by 180-day survival, faceted by shock status (Wilcoxon rank-sum, unadjusted p shown). (E) Paired day-0 and day-7 IFNɣ, IL-22, IL-17A, and IL-9 responses to anti-CD3/CD28, faceted by shock status and colored by 180-day survival. Paired Wilcoxon signed-rank test. Significance defined as *** FDR < 0.05, ** FDR < 0.1, * FDR < 0.25. (F) Paired day-0 and day-7 IL-2, IL-22, IL-17A to heat-killed *C. albicans*, faceted by shock status and colored by 180-day survival. Paired Wilcoxon signed-rank test. Significance defined as *** FDR < 0.05, ** FDR < 0.1, * FDR < 0.25.

Several day-0 responses to pan-T cell activation distinguished infected patients with sepsis from those without. Comparing infected non-septic (n = 8) and infected septic patients (n = 59), anti-CD3/CD28-induced IFNɣ was 6.7-fold lower in the septic group (median 489 [IQR 177-1,444] versus 3,271 [IQR 502-8,980] pg/mL; unadjusted p = 0.036; FDR = 0.08; **Figure 4B, Table S4**). IL-22, IL-17F, IL-17A, IL-9, IL-6 and IL-10 induction by anti-CD3/CD28 were likewise 3.9-to 22-fold lower in septic patients (all unadjusted p < 0.05; FDR < 0.1), with IL-2 and TNF-α trending in the same direction (p = 0.087 and 0.13). In addition, specific responses to heat-killed *Candida albicans* also distinguished the two groups, including higher IL-6 and IL-10 induction in infected non-septic patients (11,153 [2,397-19,847] versus 298 [76-2,191] pg/mL, p = 0.023, FDR = 0.1; and 100 [50-141] versus 6 [1-32] pg/mL, p = 0.043, FDR = 0.1; **Figure 4C**), with TNF-α 14-fold higher (p = 0.059, FDR = 0.12; **Figure S5B**). T-cell cytokine responses to HKCA (IFNɣ, IL-13, IL-17A) reached significance but were near the lower limit of detection in both groups, while IL-2, IL-22, IL-17F, IL-9, IL-4 and IL-5 were not significantly different. Together, these results indicate that among infected and hospitalized patients, sepsis is associated with a select deficit in multiple T cell associated cytokines, as could be expected given the common lymphopenia associated with sepsis, as well as a reduction in innate responses to fungal but not bacterial or TLR stimulation. Beyond these differences, septic patients showed a high degree of heterogeneity, and we next focused on understanding whether these responses to stimuli were associated with particular physiological states or outcomes.

Next, to determine whether cytokine responses were associated with clinical trajectory, and building on the comparison groups we used for flow cytometry results, we stratified day-0 responses by shock status and 180-day survival. Among patients without shock, IFNɣ responses at day-0 trended higher in survivors than in non-survivors (852 [357-3,896] vs 414 [149-552] pg/ml, p = 0.078; **Figure 4D; Table S4c**). In contrast, in septic shock, the two groups had comparably suppressed IFNɣ induction (381 [125-601] vs 486 [116-2020] pg/ml; p = 0.43).

We then focused on the subset of infected septic patients with paired day-0 and day-7 samples in both assays (n = 30). In the non-shock stratum, T-cell derived responses (IFNɣ, IL-22, IL-17A, and IL-9) in survivors showed significant improvement by day 7 (FDR < 0.25, paired comparisons across 79 stimulus-cytokine pairs in each stratum; **Figure 4E, Table S4b**); however, no significant functional recovery was observed in non-survivors without shock, nor in either survivors or non-survivors in the shock stratum. Similarly, HKCA induced responses, including IL-2, IL-22, and IL-17A, were significantly increased by day 7 in non-shock survivors (FDR < 0.25), while no significant differences were seen in non-survivors without shock, as well as in either survival groups in shock (**Figure 4F**).

Because the compositional and functional assays were performed on the same patients, we then asked whether differences in IFNɣ responsiveness reflected differences in the compartment that produces it. In the non-shock stratum at day 0, anti-CD3/CD28-induced IFNɣ correlated with memory CD4 abundance (Spearman rho = 0.60, p < 0.001; **Figure S5C-D**) and with both CXCR3+ metaclusters (Cluster 8 rho = 0.66, p < 0.001; Cluster 9 rho = 0.62, p < 0.001; **Figure S5E-F**), each expressed as a percentage of live PBMC. In the septic shock stratum, this architecture was not preserved. At day 0, anti-CD3/CD28-induced IFNɣ was uncoupled from the abundance of both CXCR3+ metaclusters (Cluster 8 rho = 0.22, p = 0.27; Cluster 9 rho = 0.10, p = 0.63). Spearman correlation networks of Tier 2 subset abundance and cytokine responses (edges filtered at |rho|≥0.4 and FDR < 0.1) demonstrated a similar relationship. In sepsis without shock the network resolved into two axes: T cell-derived cytokines, including IFNɣ, IL-2, IL-17A, IL-17F, IL-22, IL-9, correlated with memory CD4+ T cell abundance; innate cytokines, including IL-6, IL-10, TNF-α induced by whole microbes, IL-1b or TLR ligands correlated with classical monocyte abundance. The septic shock network was markedly sparser (7 versus 28 edges) and rewired, with the memory CD4-IFNɣ edge falling below FDR threshold (rho = 0.45). Instead, memory CD4 correlated with IL-5, CD45RA+CD45RO+ intermediate CD4+ T cells correlated with IL-4, IL-13 and IL-9, and innate responses tracked intermediate rather than classical monocytes (**Figure S5C, Table S4c**).

In summary, functional immune profiling demonstrated that early defective responses to T cell activation, especially blunted IFNɣ induction, as well as reduced innate responsiveness to fungal stimulation, may help to distinguish sepsis from uncomplicated infection. Within sepsis, day-0 IFNɣ induction trended with 180-day survival in patients without shock but not in septic shock. Furthermore, IFNɣ functional capacity scaled with the size of the memory CD4 compartment, with significant functional recovery observed over the first week in non-shock survivors alone. In contrast, while the memory CD4 and CXCR3+ compartments were present, they no longer tracked with IFNɣ capacity in septic shock.

### Antibody profiling links prior CMV exposure to CXCR3+ memory CD4+ T cell

In addition to examining the circulating immune cell composition and functional responsiveness, we investigated how prior exposures shape the host immune state. To comprehensively map prior microbial exposures and the range of antibody responses, we employed phage immunoprecipitation sequencing (PhIP-Seq)^22,23^. The phage display library was constructed by tiling 56-amino acid peptides across proteins from four categories **(Figure 5A)**: (1) the complete human virome (using UniRef90 representatives, with UniRef50 applied to HIV-1 to account for the overrepresentation of HIV-1 variants in UniProt); (2) bacterial epitopes with at least one epitope indexed in the Immune Epitope Database (IEDB); (3) human extracellular and secreted proteins ^24^; and (4) proteins collected from the Allergome database. Including additional negative controls, the complete library comprised 229,814 unique oligonucleotides (**Methods**) and uniform representation was validated by sequencing the produced phage pool.

**Figure 5.**
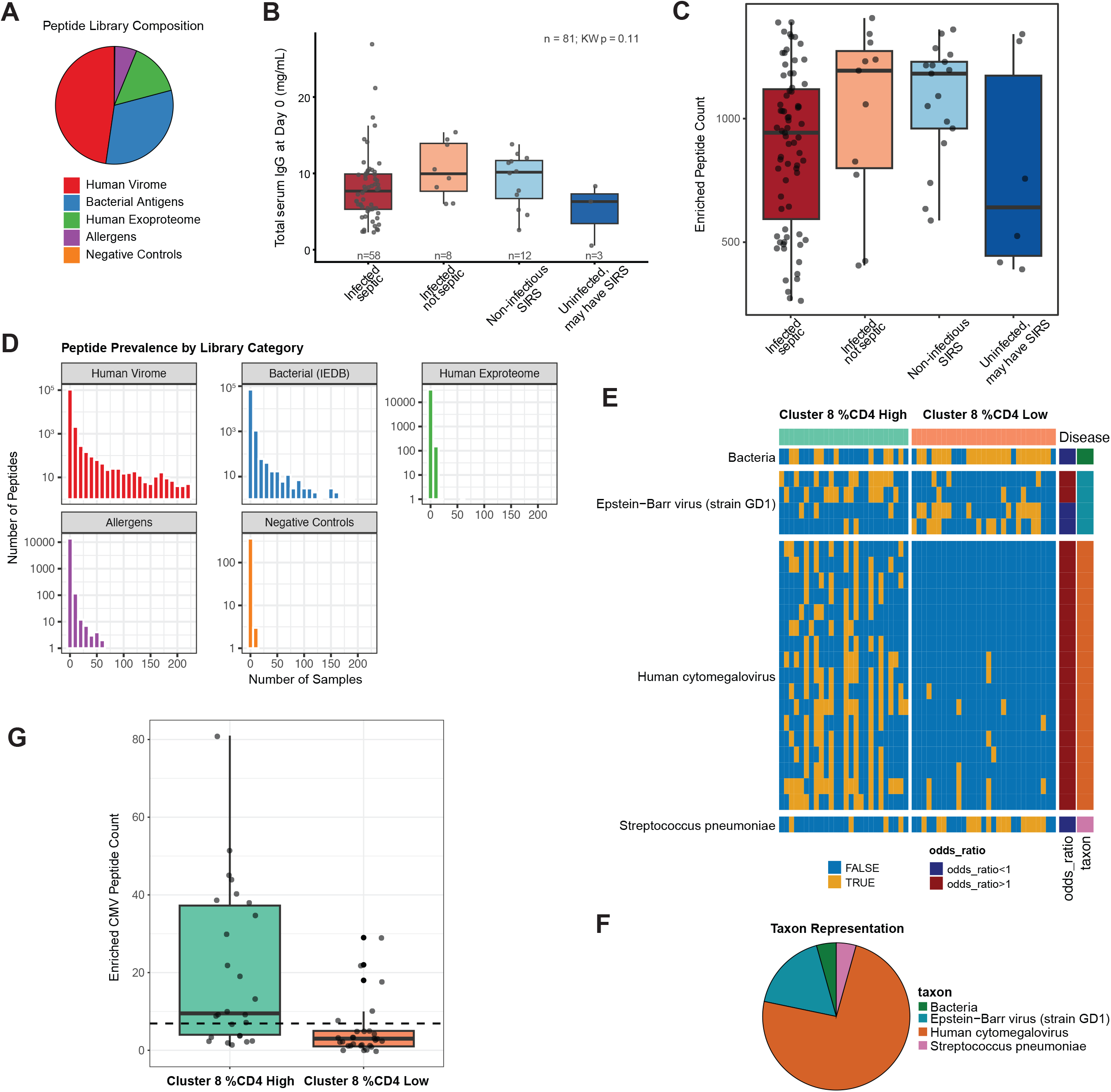
Serum antibody repertoire analysis links CMV seropositivity to immune profiles. (A) Pie chart of phage display peptide library composition. (B) Total serum IgG at day 0 by adjudicated category (infected septic n = 58, infected but not septic n = 8, non-infectious SIRS n = 12, uninfected may have SIRS n = 3). (C) Number of enriched peptides detected per sample at day 0, by adjudicated category. (D) Peptide prevalence across the cohort, faceted by library category (human virome, bacterial antigens from IEDB, human exoproteome, allergens, and negative controls). (E) Differential peptide enrichment between patients with high and low day-0 Th1 (CXCR3+, Cluster 8) abundance, split at the median abundance as a percentage of memory CD4. Each peptide tested by Fisher’s exact test with Benjamini-Hochberg correction. Each row per peptide and one column per sample, yellow indicates enrichment (FDR < 0.25). Rows are annotated by source taxon and by the direction of the association as odds ratio > or < 1. (F) Taxon representation among the 23 differentially enriched peptides in (E). (G) Boxplot of enriched CMV peptide counts between above-and below-median Th1 abundances. Median and interquartile range per group as indicated. Dotted line at 6.9 represents 5% of public epitopes indicating CMV seropositivity.

We used this library to characterize the antibody repertoire of our sepsis-at-risk cohort. The total IgG concentrations were first quantified via ELISA (**Figure 5B**) and normalized across all samples before phage immunoprecipitation. Subsequently, precipitated phages were lysed, amplified and sequenced, and significantly enriched peptides were called by detection of outliers relative to similarly represented peptides. Across the entire cohort, we detected a median of 979 (IQR 644–1216) enriched peptides per subject at day 0. The number of enriched peptides were similar between adjudicated patient categories and between day 0 and day 7 (**Figure 5C**, **Figure S6A, Table S5, Table S5b-c**). The global distribution of epitope recognition was consistent with detection of many public viral and bacterial epitopes, as expected for immunodominant epitopes across the population (**Figure 5D**). In contrast, responses targeting the human exoproteome and allergens were restricted to a small fraction of individuals and, as expected, negative controls were almost completely undetected (**Figure 5D**).

We next focused specifically on potential differences across infected septic patients at day 0 to identify previous exposures associated with specific immune or disease features. The total number of enriched peptides was similar between shock and non-shock groups (**Figure S6B**). Given the difference in CXCR3+ Th1 abundance between survivors and non-survivors of sepsis without shock, as well as the role of these cells in SOFA improvement in the shock subset, we first asked whether past exposures were associated with the frequency of this subset. Among infected septic patients with paired day 0 serum and PBMC samples (n = 55), we dichotomized patients at the median CXCR3+ Th1 (Cluster 8) abundance, expressed as a percentage of memory CD4+ T cells, and compared differential peptide enrichment between the groups by Fisher’s exact test with multiple testing correction. We focused on public responses by restricting our analysis to peptides detected in at least 20% of either group, for a total of 413 peptides tested. Twenty-three peptides were found to be differentially enriched (FDR < 0.25) (**Figure 5E-F, Table S5d-e**). Seventeen of these 23 were derived from CMV, and all were enriched in Th1-high patients, suggesting that increased responses to CMV were associated with higher CXCR3+ Th1 abundance at enrollment. To define whether this association was driven by previous exposure or a difference in the magnitude in anti-CMV responses, we next calculated the number of positive CMV peptides per subject (**Figure 5G**). Using a 5% cut-off (i.e. 6.9 peptides out of 138 public CMV peptides detected in at least 2.5% of samples, see **Methods**) to define individuals as seropositive for CMV, we observed a significant difference in exposure when separating individuals based on the frequency of cluster 8 as a percentage of memory CD4 T cells (18 positive vs 8 negative in Th1-high group, 5 vs 24 in Th1-low group, Fisher’s exact test p = 0.0001). While we were not powered to define the effect of previous exposures on survival, we note that CMV seropositivity showed a trend towards an association with survival in the group without shock (12 positive vs 9 negative in survivors, 2 vs 7 in deceased, p = 0.12) that was absent in the group with shock (5 positive vs 10 negative in survivors, 4 vs 6 in deceased, p = 1), which is consistent with the observed association between Th1 cells and survival in sepsis without shock. Similarly, in our Kaplan-Meier analysis, CMV seropositivity showed a trend toward an association with survival in sepsis without shock, but not in septic shock (log-rank p = 0.093 in sepsis without shock, p = 0.758 in septic shock; **Figure S6C, TableS5f**).

Together, these data indicate that prior CMV exposure is associated with a higher frequency of Th1 cells early during sepsis and provides additional insights into the factors that drive the heterogeneity of these populations and subsequent sepsis outcomes.

## Discussion

In this study, we defined the immune states of 89 hospitalized adult patients at risk for sepsis, using a suite of three complementary assays: spectral flow cytometry of circulating immune composition, *ex vivo* functional stimulation with cytokine quantification, and antibody repertoire profiling by PhIP-seq. We demonstrate that within the memory CD4+ compartment, a CXCR3+CCR4-Th1-polarized cluster at enrollment was enriched in 6-month survivors of sepsis without shock, tracked recovery of organ dysfunction in the first week of septic shock, and was coupled to the magnitude of T cell IFNɣ production after restimulation. Importantly, while shock is associated with poor survival, individuals without shock but with low CXCR3+ frequencies at time of enrollment had similar outcomes to individuals with shock. Additionally, seromic analyses demonstrated a strong association between high frequencies of CXCR3+ memory CD4 T cells and previous CMV exposure, suggesting that this virus is one of the factors influencing the level of these cells in the context of sepsis. However, the physiological state of the host altered the prognostic role of this immune profile. Specifically, the presence of septic shock uncoupled Th1 abundance from IFNɣ capacity and abolished its association with survival. While previous sepsis endotyping efforts have predominantly stratified patients based on blood transcriptomic signatures, our findings suggest a different framing: the value of prognostic immune signatures can be modulated by the physiologic state of the host, and precision immune biomarkers in sepsis should therefore be interpreted in the physiologic context, such as whether shock is present.

Our finding that a Th1-polarized memory CD4+ compartment and preserved IFNɣ production are associated with survival in sepsis without shock aligns with the growing recognition that a principal driver of late sepsis mortality is a state of immune paralysis ^25^. In sepsis-induced immunosuppression, the depletion and exhaustion of the T cell compartment impair clearance of the primary infection and leave patients susceptible to secondary infections and viral reactivation ^26–28^. The major cytokine produced by Th1 cells is IFNɣ, which plays a critical role in host defense against viral and intracellular pathogens. Specifically, IFNɣ upregulates antigen presentation capabilities, activates macrophage microbicidal effector functions, and induces a potent cellular antiviral state ^29^. The key immunological role of IFNɣ is the biological basis for leveraging recombinant human IFNɣ for the treatment of sepsis, as recently evaluated in the multicenter, randomized placebo-controlled precision immunotherapy clinical trial, ImmunoSep^8^. In this trial, blood ferritin concentrations and the expression of HLA-DR on monocytes were used to stratify levels of immune activation, with patients randomized to receive targeted therapies for either hyperinflammation or sepsis-induced immunoparalysis. Our work complements this randomized clinical trial observation but differs in several meaningful ways: first, while IFNɣ administration improved organ dysfunction trajectories in this trial, it did not alter survival; in our cohort high frequencies of Th1 cells at day of enrollment directly tracked with 6-month survival in sepsis without shock. This may reflect differences in the populations studied but also the effect of systemic administration compared to dynamic changes in Th1 populations. Defining the specificity and functional role of these cells will be essential to translate this observation into potential interventions aimed at longer survival. We note that in most of these patients, circulating Th1 frequencies were higher, and more predictive, at day 0 than day 7, suggesting that this compartment is dynamic during this window of acute inflammation.

However, our findings suggest that this specific immune axis targeted by these precision therapies may be fundamentally rewired by the presence of shock. Interestingly, Th1 cells were still associated with improved organ dysfunction trajectories in the first week of septic shock, but not with improved six month survival. This could be explained by the increased severity of shock or other comorbidities and factors influencing care decisions in this group, or by functional changes in the CXCR3+ populations. In our cohort, instead of being robustly correlated with T-cell derived IFNɣ, CD4+ memory and intermediate T cells in septic shock were instead more strongly correlated with Th2 cytokine production. Indeed, prior studies in models of polymicrobial sepsis have demonstrated that CD4+ T cells shift from a pro-inflammatory Th1 phenotype to an anti-inflammatory Th2 profile ^30^. This phenomenon has also been observed in human cohorts and associated with 28-day survival ^31^.

Interestingly, as septic shock leads to metabolic abnormalities that profoundly increase mortality ^1^, it likely also directly influences T cell function. A prior study demonstrated that conventional CD4+ memory T cells, including Th1 cells, lack the metabolic plasticity to augment glycolytic capacity under critical illness-related metabolic stress, leading to cellular attrition and functional impairment ^32^. Similarly, extracellular lactic acid present in tumors was shown to have a suppressive effect on CD4+ T cell function and impaired IFNɣ production ^33^. Therefore, we hypothesize that the metabolic milieu of septic shock restricts CD4+ T cell metabolism and normal effector function, driving the functional decoupling we observe. Further mechanistic studies are required to establish whether reversing these metabolic impairments can restore Th1 effector function and improve survival in septic shock.

To define how immune history may impact sepsis course and immune parameters, we also employed antibody profiling by PhIP-Seq to evaluate how prior exposure may impact immune profiles, and found that CMV exposure was associated with above-median CXCR3+ Th1 abundance. Our findings are consistent with a prior twin study demonstrating that CMV serostatus strongly influenced immune parameters ^16^ and an immunotherapy study showing improved outcomes and reduced immune adverse events in CMV-exposed subjects undergoing checkpoint blockade therapy for melanoma ^34^. CMV reactivation can occur during sepsis ^35^, likely due to perturbed immunosurveillance. The association between reactivation and outcomes has been debated ^36,37^, but a recent study has associated CMV-seropositivity with worse outcomes and distinct immune predictors ^38^, without stratifying patients by shock status. While we did not detect a significant effect on survival, we observed a trend towards improved outcomes in sepsis without shock, likely driven by the relationship between Th1 cells and survival on one hand and Th1 cells and CMV on the other. Larger studies stratifying by both shock and CMV serostatus will be required to understand how these two parameters influence survival.

We acknowledge several important limitations. First, while our sample size (n = 89) is robust for *ex vivo* functional assays and multiparameter flow cytometry, it is modest compared to large-scale bulk transcriptomic cohorts; stratification by shock further constrained our sample size.

Consequently, the number of outcome events limited our ability for multivariable modeling, although our primary findings remained stable in sensitivity analyses. A larger, multicenter validation cohort would be useful to confirm our findings to resolve finer-grained outcomes such as susceptibility to recurrent infection. Second, our longitudinal analysis at day 7 is subject to informative censoring, as both early discharges and deaths preclude paired sampling.

Therefore, our observations regarding expansion of the memory T cell compartment and functional IFNɣ recovery at day 7 are restricted to patients who survived and remained hospitalized, requiring careful interpretation regarding survivorship bias. Third, our analysis is confined to the peripheral blood compartment. While circulating PBMCs are the most clinically accessible sample for real-time biomarker assessment during critical illness, they may not reflect the immune environment at sites of primary infection as well as organ injury, and defining the role of Th1 cells at these sites will be essential to further define their role in protection.

In conclusion, this work demonstrates the value of combining multimodal immune profiling with stratification by shock status in resolving sepsis heterogeneity and in designing or selecting immune-focused interventions, for example aimed at improving outcomes in patients without shock but with low Th1 abundance and poor outcomes.

## Supporting information

Supplementary Tables (combined)

## Acknowledgments

We thank the patients who contributed to this study and their families. This work was supported by the Center for Integrated Solutions for Infectious Diseases (to L.C, A.E.W, J.D.). R.J.X. is a cofounder of Jnana Therapeutics and Convergence Bio, a scientific advisory board member at Nestlé, Magnet BioMedicine and Arena BioWorks, and board director at MoonLake Immunotherapeutics; these organizations had no role in this study. The other authors declare no competing interests.

## Methods

### Study cohort and sample processing

#### Study design and participants

We prospectively enrolled adult patients admitted to the Brigham and Women’s Hospital, a tertiary academic medical center in Boston, Massachusetts, enrolling adults with suspected sepsis from the ED, inpatient wards, and ICUs under the IRB protocol #2023P000161 (Sample collection and clinical assessment of individuals with or at increased risk for infectious diseases). Eligible patients included adults (≥18 years) with suspected infection, defined as a new antibiotic administered, together with a microbiological culture or PCR obtained within the previous 24 hours and at least two Systemic Inflammatory Response Syndrome (SIRS) criteria. Exclusions were limited to contraindications to biospecimen collection as determined by the study team or clinical team. Patients were identified through a three-layer screening process performed on weekdays that included (1) automated electronic medical-record screening, operationalized as daily automated electronic health record reports (Epic, Epic Systems Corporation, Verona, Wisconsin) spanning the ED and all inpatient locations with a 24-hour lookback for two or more signs of SIRS, microbiologic culture or PCR, and receipt of a new antibiotic; (2) a sepsis enrichment step whereby trained non-physician study staff performed structured chart review (estimated 3-5 minutes per chart) for evidence of positive microbiologic cultures or PCR results, chart documentation of confirmed or high clinical suspicion of sepsis by ICU or infectious disease consultants; and/or (3) in-person discussion with the primary clinical team. Individuals who passed all three steps were flagged for enrollment. After enrollment, whole blood was collected on days 0 and 7 (+/-1 day) post-enrollment, in patients who were alive and still admitted. All patients with at least one index sample in the first seven days were considered enrolled and retained for clinical-outcome analysis. Clinical data were abstracted from the electronic health record into REDCap electronic data capture tools hosted at Brigham and Women’s Hospital and included demographics, comorbidities, medications, microbiology, and organ-dysfunction measures. Daily Sequential Organ Failure Assessment (SOFA) scores were calculated from clinical and laboratory data extracted from the Mass General Brigham Snowflake data warehouse. For each patient-day, the six organ-system components (respiratory, cardiovascular, hepatic, coagulation, renal, and neurological) were each scored from the most abnormal value recorded that day, yielding the daily maximum SOFA score as the sum of the six components. When the clinical or laboratory data required for a component were unavailable on a given day, that component was assumed to be normal and assigned a score of 0. Daily scores were restricted to the index hospitalization, which ended at discharge or at death or transition to comfort-measures-only (CMO) care.

Because withdrawal of vasopressor and ventilatory support can lower SOFA scores independently of organ recovery, we replaced the SOFA score recorded on the day of death or transition to CMO with the preceding day’s score to avoid misclassifying treatment withdrawal as clinical improvement. Outcomes included index hospitalization length of stay, ICU admission and ICU days, occurrence of shock, in-hospital and all-cause 30-, 90-, and 180-day mortality, and 30-and 90-day readmission.

#### Clinical Adjudication and Definitions

Enrolled patients underwent structured physician adjudication at least 14 days after enrollment to assess the clinical course, microbiological results, imaging, and treatment response. Two independent physician adjudicators applied a predefined rubric to determine the presence or absence of infection, the presence or absence of sepsis by Sepsis-3 criteria, the presence or absence of septic shock, and the anatomic and microbial source of infection. Septic shock was identified based on Sepsis-3 criteria: vasopressor requirement to maintain a mean arterial pressure of 65 mm Hg or greater and serum lactate level greater than 2 mmol/L (>18 mg/dL) in the absence of hypovolemia ^1^.

Discordant results were reviewed by a third physician tie-breaker. Each patient was assigned to one of four categories: (1) infected septic, (2) infected but not septic, (3) undifferentiated SIRS, (4) and non-infectious SIRS. Infectious pathogen and anatomical sources were adjudicated using relevant microbiological, clinical, and imaging data.

#### PBMC and Serum isolation

Whole blood was collected into serum separator tubes and EDTA blood collection tubes via standard phlebotomy procedures and processed within the same day of collection. To isolate serum, the serum separator tubes were centrifuged at 1,000 x g for 5 minutes at 4°C with maximum acceleration and brake. Isolated serum was aliquoted into cryogenic tubes and immediately frozen at-80°C. For peripheral blood mononuclear cell (PBMC) isolation, room temperature EDTA-anticoagulated blood was diluted 1:1 with PBS and layered onto 14 mL of Ficoll-Paque Plus (density 1.077 g/mL, GE Healthcare, Sweden)) in SepMate (STEMCELL) tubes. Density gradient centrifugation was performed at 1,200 x g for 10 minutes at 20°C with maximum acceleration and brake. Following centrifugation, 2 mL of the plasma layer was removed, the remaining plasma and PBMC layer was poured into a new 50 mL conical tube, diluted 1:1 with cold RPMI-1640 supplemented with 2% FBS, and centrifuged at 350 x g for 5 minutes at 4°C for the first wash. The cell pellet was resuspended in 1-2 mL of ACK lysis buffer and incubated for 1 minute to remove red blood cell contamination and washed again with cold RPMI + 2% FBS. An aliquot of the cell suspension was diluted with trypan blue for cell counting with hemocytometer. Finally, the washed PBMC pellet was resuspended in cold CryoStor CS10 (Stemcell Technologies) at 5 x 10⁶ cells/mL in cryotubes and incubated at 4°C for 10 minutes. The cryotubes were then transferred to controlled-rate cooling containers and immediately placed in a-80°C freezer for a minimum of 12 hours and a maximum of 90 days.

Subsequently, samples were transferred on dry ice to the vapor phase of liquid nitrogen for long-term storage.

#### Multiparameter Spectral Flow Cytometry

Cryopreserved PBMCs were rapidly thawed in a 37°C water bath and resuspended in RPMI-1640 containing HEPES and L-glutamine, supplemented with 10% (v/v) FBS, 1X MEM non-essential amino acids (0.1 mM each), 1 mM sodium pyruvate, 55 µM 2-mercaptoethanol, 100 U/mL penicillin, and 100 µg/mL streptomycin (complete media). Cells were quantified using an automated cell counter, and 5 x 10^5^ to 1 x 10^6^ cells per donor underwent antibody staining for flow cytometry. The remaining cells were used for *ex vivo* stimulation (see below). Cells were first stained with Live/Dead Fixable Near IR viability dye (ThermoFisher, Cat no. L34981), according to the manufacturers’ instructions, for 30 minutes at 4℃. After washing with FACS (1x PBS + 2% fetal bovine serum [FBS] + 1 mM EDTA) buffer, cells were incubated with Human TruStain FcX (BioLegend, Cat no. 422302) for 20 minutes at 4℃. Two antibody panels were used to characterize immune cell composition: Panel A (T-cell subsets) consisted of 12 fluorochrome-conjugated antibodies targeting CD25, CCR4, CXCR3, CCR10, CD3, CD4, CD8, CD45RO, PD-1, CD127, CD45RA, and CCR6. Panel B (Myeloid and B-cell populations) included 10 antibodies targeting HLA-DR, CD11c, CD40, CD206, CD19, CD3, CD14, CD16, CD5, CD56, and CD123 (see **Methods**). Antibody staining for both panels was performed for 20 minutes at 4℃. Data acquisition for all samples was performed using a Sony ID7000 Spectral Analyzer. Single-color compensation controls were prepared using UltraComp eBeads (Thermo Fisher) for all fluorochromes aside from the Live/Dead IR single-color, which was performed on PBMCs heat-shocked for 5 minutes at 42℃. Spectral unmixing was done using the Sony ID7000 software unmixing algorithm after autofluorescence subtraction defined with unstained donor PBMCs. Unmixed FCS files were exported for downstream analysis in FlowJo v10 (BD Biosciences) and gated as shown in Figure S1A to define population frequencies.

#### *Ex Vivo* PBMC Stimulation and Cytokine Quantification

Cryopreserved PBMCs were thawed, washed, and resuspended in complete RPMI then plated in 96-well V-bottom plates at a density of 50,000 cells per well. Cells were allowed to settle at 37°C for 1 hour before stimulation with the following ligands: heat-killed *Candida albicans* (HKCA), *Staphylococcus aureus* (HKSA), and *Escherichia coli* (HKEB) at 10⁷ units/mL; lipopolysaccharide (LPS) at 500 ng/mL; R848 at 1µM; recombinant human IL-1β at 25 ng/mL; and human T-activator anti-CD3/anti-CD28 Dynabeads at a 1:1 bead-to-cell ratio (ThermoFisher; Cat no. 11161D, or complete media alone as a control. The final volume for cell suspension plus stimuli was 50µL per well (1M cells/mL final). After stimulation for 16 hours at 37°C, plates were centrifuged at 1,000 g for 10 minutes to clear cellular debris, and supernatants were harvested and immediately stored at-80°C. Cytokine concentrations in the supernatants were quantified using the LEGENDplex Human Th Cytokine Panel (12-plex; BioLegend), which includes IL-2, IL-4, IL-5, IL-6, IL-9, IL-10, IL-13, IL-17A, IL-17F, IL-21, IL-22, IFN-*γ*, and TNF-*α*. The assay was performed according to the manufacturer’s instructions with a modified reagent volume of 12.5 µL of capture beads, detection antibodies, and Streptavidin-PE (SA-PE) per well. Bead fluorescence was acquired on a CytoFLEX S flow cytometer (Beckman Coulter) and analyzed using the LEGENDplex Data Analysis Software (BioLegend).

### Serum antibody profiling

#### Construction of epitope set and generation of phage library

The library contains peptides from the complete human virome (∼48%), bacterial immunodominant proteins (∼31%), human cytokines and exoproteome (15%), food allergens (6%) and negative controls.

#### Human Virome

We downloaded all full-length protein sequences from viruses with human host present in the UniProt database (March 2023) and collapsed them at 90% identity by using Uniref90 clustering algorithm (except for HIV-1). The algorithm represents each Uniref90 cluster of proteins with at least 90% similarity with a single representative sequence, and we obtained 10,133 Uniref90 representative sequences. Because of the high number and variability of HIV1 proteins in Uniprot, we clustered them at 50% identity by using Uniref50 clustering algorithm, resulting in 894 Uniref50 HIV-1 protein clusters.

#### Bacterial Immunodominant Proteins

For bacterial immunodominant proteins, we downloaded from IEDB (March 2023) all protein sequences from bacterial pathogens and commensals with at least one known epitope (B cell or T cell epitope), collapsed at 90% identity by the Uniref90 clustering algorithm, resulting in 4,444 bacterial protein clusters (including 34 protein sequences present in IEDB that were not present in Uniprot, could not be clustered by Uniref90).

#### Food and other Allergens

The protein sequences of allergens present in Allergome database (June 2024), excluding human, mouse, rat and arthropods, were collapsed at 90% identity by the Uniref90 clustering algorithm, resulting in 1,506 Uniref90 protein clusters.

#### Human Cytokines and Exoproteome

The protein sequences of human cytokines and chemokines present in KEGG database (March 2023) were collapsed at 90% identity by the Uniref90 clustering algorithm, resulting in 148 protein clusters. A comprehensive list of 3,394 protein sequences from human secreted proteins and extracellular domains of surface proteins, described in a recent exoproteome-microbiome interaction study ^39^, were included in the library.

#### Biological and Technical Negative Controls

The protein sequences of 38 abundant human proteins (such as serum albumin, histones and ribosomal proteins) that should not elicit an antibody response in healthy individuals were included as negative biological controls ^40^.

In addition, we included 100 56-mer peptides of random amino-acid sequences that should not be recognized by antibodies as technical negative controls ^40^.

#### Peptidome Phage library construction

The oligonucleotide library was designed following the published protocol ^23^, using the pepsyn Python package as outlined below. In short, the protein sequences from all the different library modules described above, were split into peptide tiles of 56 amino-acids with 28 amino-acid overlap, followed by removal of any peptides that were redundant (100% identity) using the CD-HIT clustering algorithm. In total, the PhIP-seq peptidome library consists of 229,814 peptides. The peptides tiles were reverse translated to oligonucleotides with an optimized *E. coli* codon usage algorithm, EcoRI and HindIII restriction cloning sites were removed by silent codon substitution and adaptor PCR-primer binding sequences were added at both ends. The resulting 206-nt oligonucleotide sequences were synthesized by pooled oligo synthesis by Agilent (SurePrint High Fidelity (HiFi) Oligo pools).

The resulting oligonucleotide library was amplified by PCR and was cloned to the EcoRI/HindIII digested T7Select-103b vector arms, followed by packaging into T7 bacteriophages (T7Select10-3 Cloning Kit, MilliporeSigma), and phage amplification in liquid bacterial cultures using the manufacturer suggested protocol.

#### Phage immunoprecipitation

Phage Immunoprecipitation was performed based on the published VirScan Bio-protocol ^41^ with slight modifications as outlined below. In short, 96-well deep well plates were blocked with TBST 3% BSA overnight. Phage-antibody complex formation was performed using 2 µg Serum IgG and 2.3 x1010 phages diluted in 1 mL Phage Extraction buffer (∼105 phages per library member ^23^) and the samples were incubated at a rotator at 4°C overnight. 40 µL of 1:1 mixtures of Protein A and Protein G magnetic beads (Thermo Fisher Scientific, 10008D and 10009D) were added and incubated with the Phage-antibody complexes for 4 hours on a rotator at RT. The beads were transferred to PCR plates and were washed 4 times with PhIP-Seq Wash Buffer followed by one wash with PhIP-Seq Wash Buffer w/o NP-40. Finally, the beads were resuspended in 40 µL of ddH2O followed by phage lysis at 95°C. Pooled DNA library construction for Illumina sequencing was performed in 3 PCR amplification steps with Q5 polymerase (New England Biolabs, M0493L) according to the manufacturer’s recommendations (20 cycles PCR1: tcgtcggcagcgtcagatgtgtataagagacagGTTACTCGAGTGCGGCCGCAAGC and gtctcgtgggctcggagatgtgtataagagacagATGCTCGGGGATCCGAATTC; 10 cycles PCR2: Illumina Nextera dual index primers; 1 cycle PCR3 (of PCR2 pools): AATGATACGGCGACCACCGA and CAAGCAGAAGACGGCATACGA). PCR3 products were cut from agarose gel and purified twice (Zymo gel extraction kit, AMPure XP beads) and sequenced on an Illumina NextSeq 2000 instrument with custom primers for R1:ttactcgagtgcggccgcaagctttca (60bp) and for R2: tgtgtataagagacagatgctcggggatccgaattct (60 bp).

#### Immunoglobulin quantification

Serum IgG concentrations were measured using the Human Immunoglobulin G (IgG) ELISA Antibody Pair Kit (Stemcell, Cat # 01994) following the manufacturer’s protocol.

#### Library sequencing

Libraries were sequenced on a NextSeq 2000 sequencer (Illumina) using custom primers and the following read configuration: i1, i2: 8bp, r1, r2: 60bp. Raw base calling files were demultiplexed into fastqs with BCL Convert v4.2.7 (Illumina).

## Data analysis

### Analysis software

All analyses were performed in R 4.5.2. Principal packages were flowCore 2.22.1 and FlowSOM 2.18.0 (cytometry data handling and clustering), ConsensusClusterPlus 1.74.0 (cluster stability), uwot 0.2.4 (UMAP), vegan 2.7.3 (PERMANOVA and PERMDISP), survival 3.8.3 and survminer 0.5.2 (survival analysis), and ggplot2 4.0.2, ggpubr 0.6.3, patchwork 1.3.2, pheatmap 1.0.13 and ComplexHeatmap 2.26.1 for visualization.

### Unbiased clustering of functional T-helper populations

Manually gated CD4+ memory T (MemCD4) populations were exported from FlowJo as individual FCS files containing the fluorescent intensities of chemokine receptors CCR4, CCR6, CCR10, and CXCR3, selected to define functional T-helper (Th) populations. These FCS files were imported into R using the flowCore package. The marker intensities were transformed using the inverse hyperbolic sine function ^42^ and z-score normalized per marker across all samples and all cells. Subsequently, unbiased clustering of functional Th populations was performed using FlowSOM ^43^, where a 10 × 10 self-organizing map (100 nodes) was trained on the four transformed markers, a minimum spanning tree was built, which was then partitioned into k = 15 metaclusters by consensus hierarchical clustering. Clusters were annotated by calculating the z-scored median fluorescence intensity (MFI) for each marker, visualized via hierarchical clustering in pheatmap (using column-wise centering and scaling, Euclidean distance, complete linkage). The abundance of each Th cluster was quantified as a relative percentage of the total MemCD4 compartment and total live PBMCs. The MemCD4 populations were downsampled to a maximum of 2,500 cells per sample and dimensionality reduction was performed via Uniform Manifold Approximation and Projection (UMAP) using the uwot package for visualization.

### Cytokine Stimulation Analysis and Correlation Networks

Raw cytokine concentrations were derived from bead fluorescence using the LEGENDplex Data Analysis Software. All subsequent data processing, statistical analysis, and visualization were performed in R 4.5.2. Induction was defined as the difference between the stimulated well and unstimulated controls, calculated per stimulus-cytokine pair for each donor, negative values were set to 0 (formula description:*cytokine induction* = *max*(0, *stimulated* − *control*)). For display on logarithmic axes, values below 1 pg/mL were set to 1 pg/mL. Group comparisons were made by Wilcoxon rank-sum test for two groups and Kruskal-Wallis for more, with Benjamini-Hochberg correction; paired day-0 to day-7 comparisons used the Wilcoxon signed-rank test.

To summarize covarying modules of composition and function, correlation networks were constructed within each shock stratum between the 19 Tier 2 sublineage frequencies and the 34 significant stimulus-cytokine pairs (defined as a mean induction greater than 10 pg/mL across the patient cohort and statistical significance [p< 0.001] by one-sample t-test of the induction values against a null of zero). Benjamini-Hochberg correction was applied within stratum. Edges were drawn where |ρ| ≥ 0.4 and FDR < 0.1, with identical, pre-specified thresholds in both strata. Networks were rendered with igraph and ggraph.

### PhIP-seq Alignment and hit calling

After quality control with fastqc-0.11.9, reads were aligned using bowtie2 to a custom library prepared with the original oligonucleotides included in the library (see above). After filtering, properly aligned reads were counted using samtools, by first extracting reads of interest (samtools-f 67) and counting them with idxstats. A custom Perl script was used to generate a long form counts table, which was transformed into a short table format in R 4.4.1.

### Differential abundance testing

Enriched peptides were determined in R using the *edgeR* function from the *beer* package, with peptides achieving a p-value < 0.05 across both technical replicates assigned as true hits. To integrate immune history with the functional T-helper response, patients were stratified into above-and below-median based on their CXCR3+ Th1 abundance within each shock stratum. To identify differentially enriched peptides between these subsets, Fisher’s exact tests were performed on peptides meeting abundance thresholds i.e. detected in at least 20% of one of the groups, with statistical significance defined as a Benjamini-Hochberg false discovery rate (FDR) < 0.25. To define CMV exposure, we summed all positive CMV peptides for each patient and, based on empirical analyses, used a threshold of 5% of public epitopes (epitopes detected in at least 2.5% of the cohort) to call CMV seropositivity.

### Survival analysis

Survival time was censored at 180 days. Survival functions were estimated by the Kaplan-Meier method with log-log transformed confidence intervals, compared by log-rank test, and modeled by Cox proportional hazards regression as indicated. The continuous Th1 abundance variable was dichotomized by splitting at the median of the relevant shock stratum, with patients at or above the median assigned to the high group; stratum-specific medians were used. Sensitivity analyses adjusted separately for age, maximum SOFA score and immunocompromised status were performed.

## Supplemental Figures

**Figure S1.**
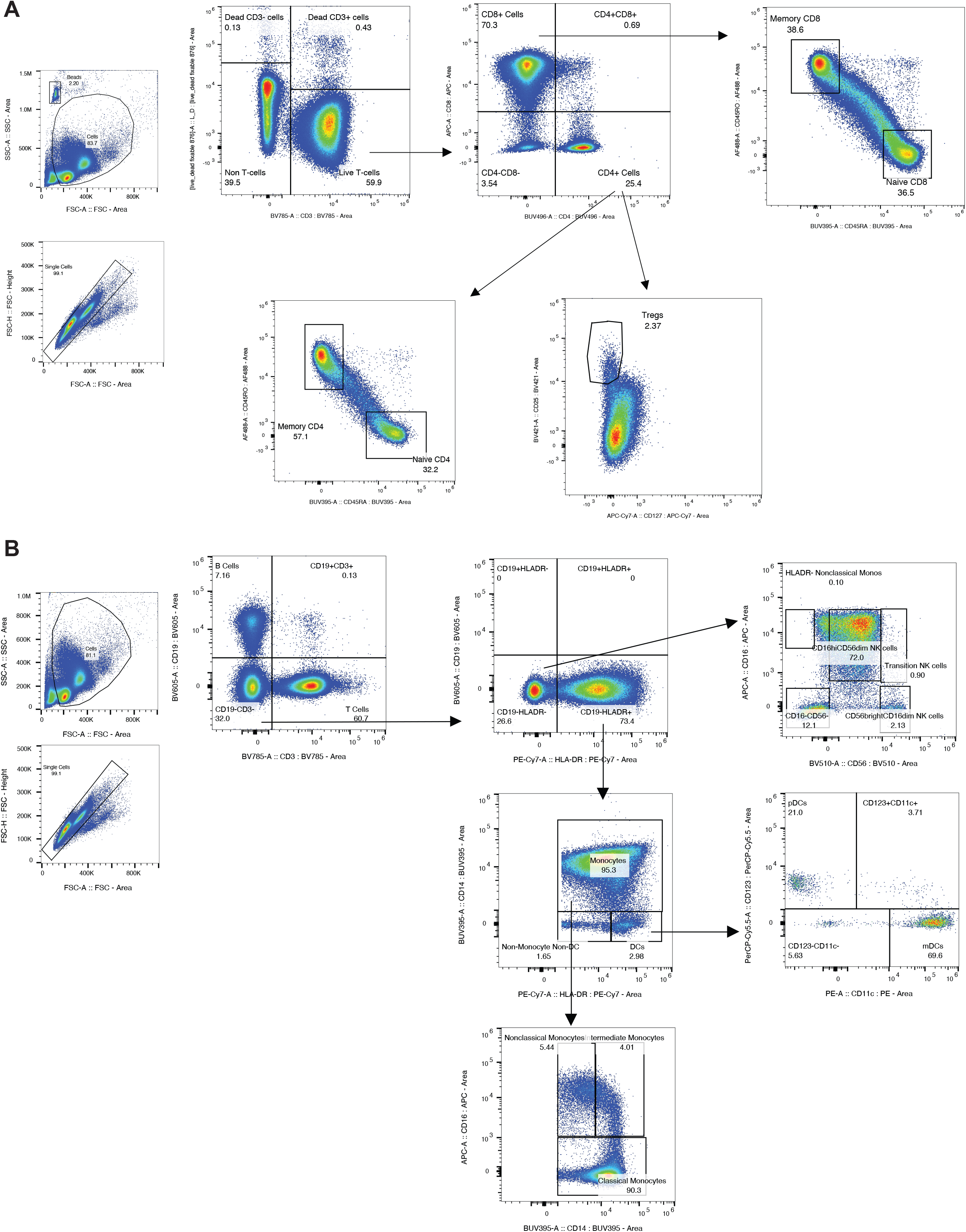
Flow cytometry gating strategy. (A-B) Gating strategy employed for the lymphoid (A) and myeloid (B) panel demonstrating the gating of Tier 1 and Tier 2 populations described in Figure 1C

**Figure S2.**
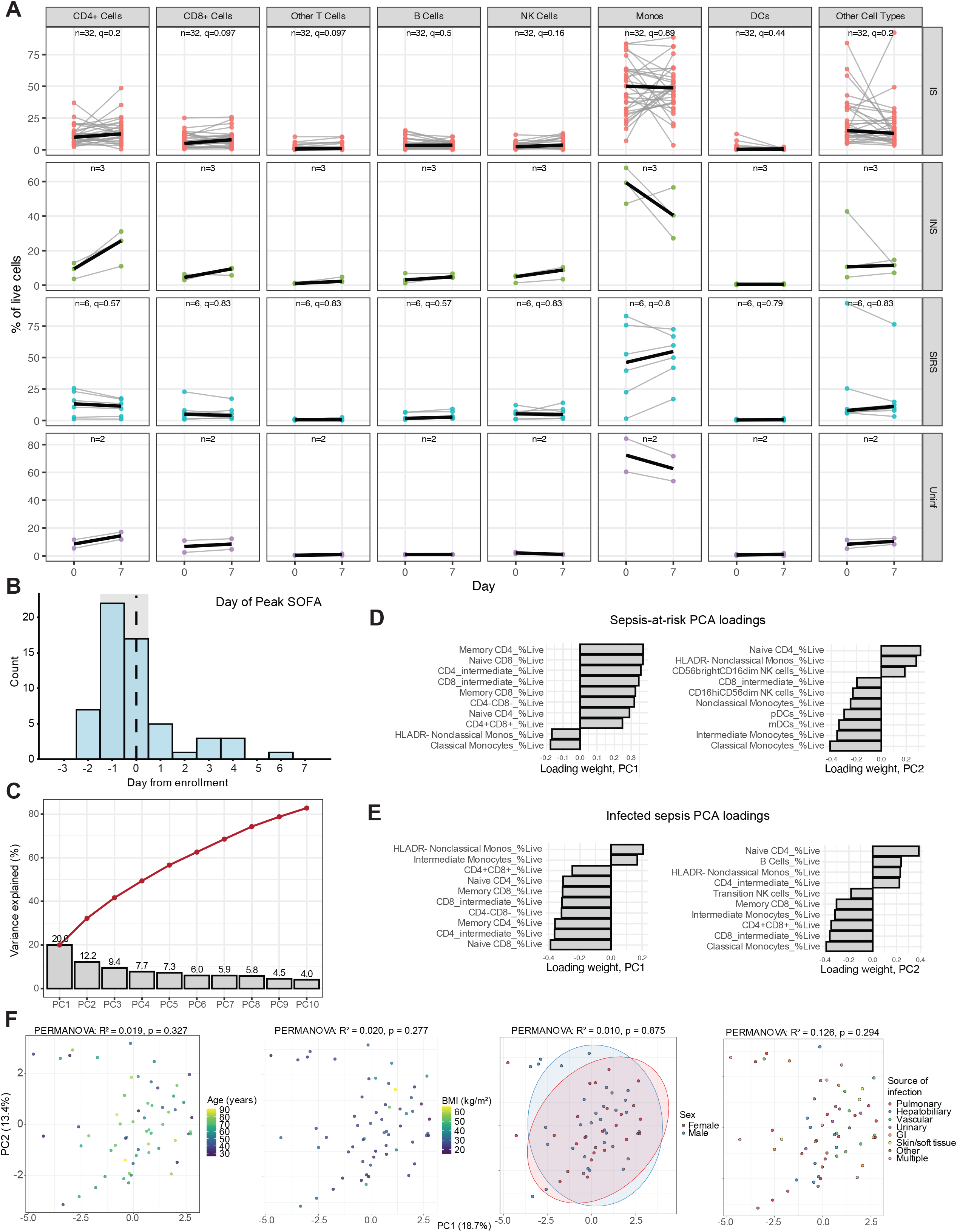
Overview of longitudinal composition and principal component analysis of composition at enrollment. (A) Paired day-0/day-7 trajectories for each Tier 1 lineage. One grey line per patient, group median in black, stratified by Tier 1 lineage (columns) and adjudicated category (rows). Paired samples:infected septic n = 32, non-infectious SIRS n = 6, infected but not septic n = 3, uninfected n = 2. Within-group Wilcoxon signed-rank tests with BH-FDR as indicated. (B) Distribution of the day of maximum SOFA score relative to enrollment among infected septic patients (n = 65). Median 0 (IQR −1 to 0, shaded). (C) Scree plot for the sepsis-at-risk PCA (Figure 1G/H). The red line indicates the cumulative total. (D) Top 10 loadings on PC1 and PC2 of the sepsis-at-risk PCA (Figure 1G/H). (E) Top loadings on PC1 and PC2 of the infected septic PCA (Figure 1I/J). (F) Day-0 infected septic PCA, colored by clinical covariates from left to right: age, body mass index, sex, and anatomic source of infection.

**Figure S3.**
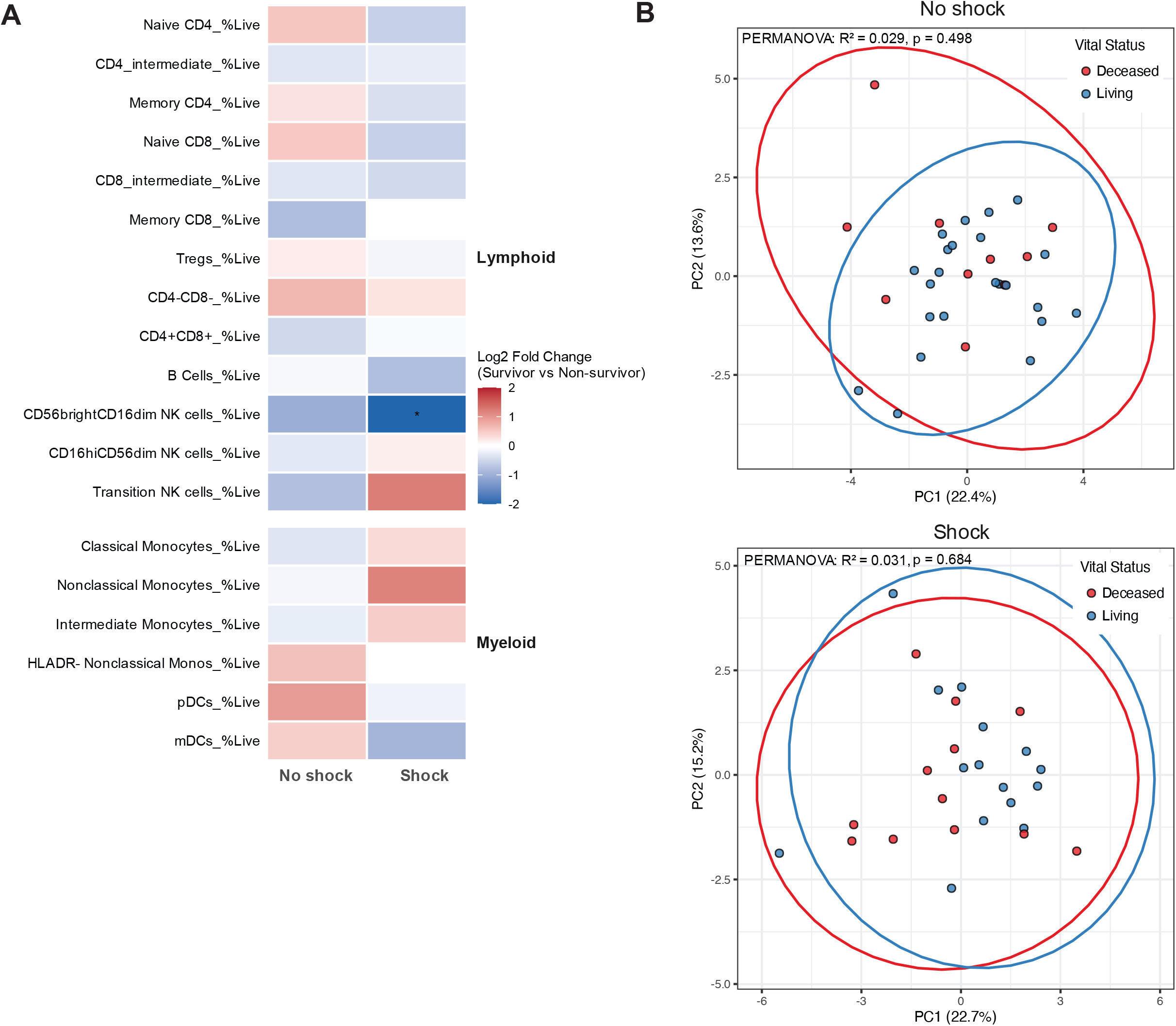
Association between PBMC composition and 180-day outcomes. (A) Tier 2 sublineage log2 fold change in day-0 abundance (% of live PBMCs), survivors versus non-survivors, within each shock stratum (no shock survivors n = 23, non-survivors n = 10; shock survivors n = 15, non-survivors n = 12). Wilcoxon rank-sum with BH-FDR across the 19 sublineages within a stratum, *, FDR < 0.25. (B) PCA of day-0 Tier 2 composition fitted separately within each shock stratum (top: sepsis without shock, bottom: septic shock), colored by 180-day vital status with 95% confidence ellipses. PERMANOVA: no shock R² = 0.029, p = 0.498; shock R² = 0.031, p = 0.684.

**Figure S4.**
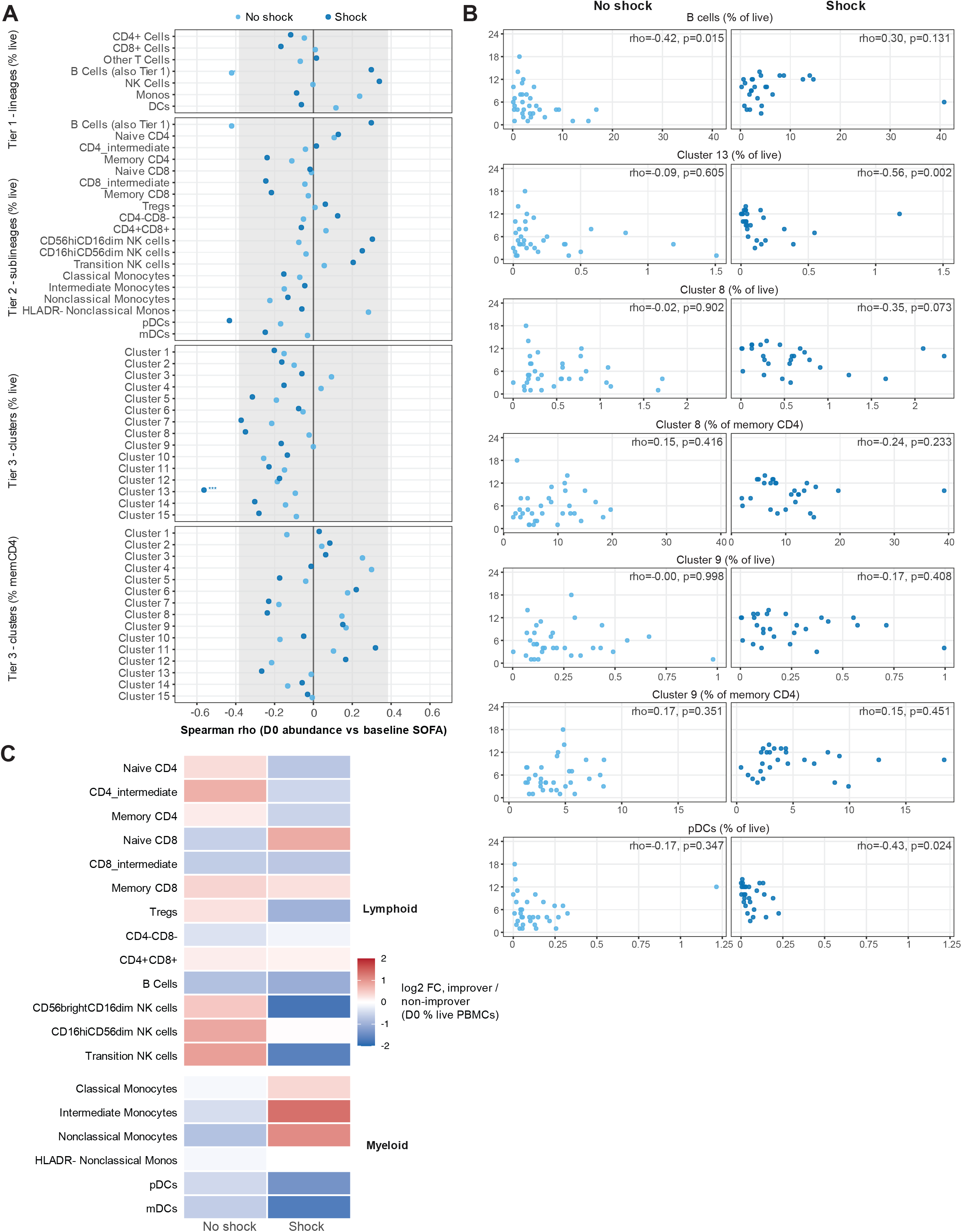
Association between PBMC composition and SOFA scores baseline and trajectories. (A) Spearman correlation between day-0 abundance and baseline (day-0) SOFA score for every population at all three tiers, by shock stratum, with BH-FDR within tier × stratum. The shaded band marks |rho| < 0.38, the magnitude required to reach p < 0.05 at given sample sizes. (B) Scatter plots of day-0 abundance against baseline SOFA for selected populations, with Spearman rho and p. (C) Tier-2 sublineage log2 fold change, improvers versus non-improvers by ImmunoSep definition. BH-FDR across 19 sublineages within stratum.

**Figure S5.**
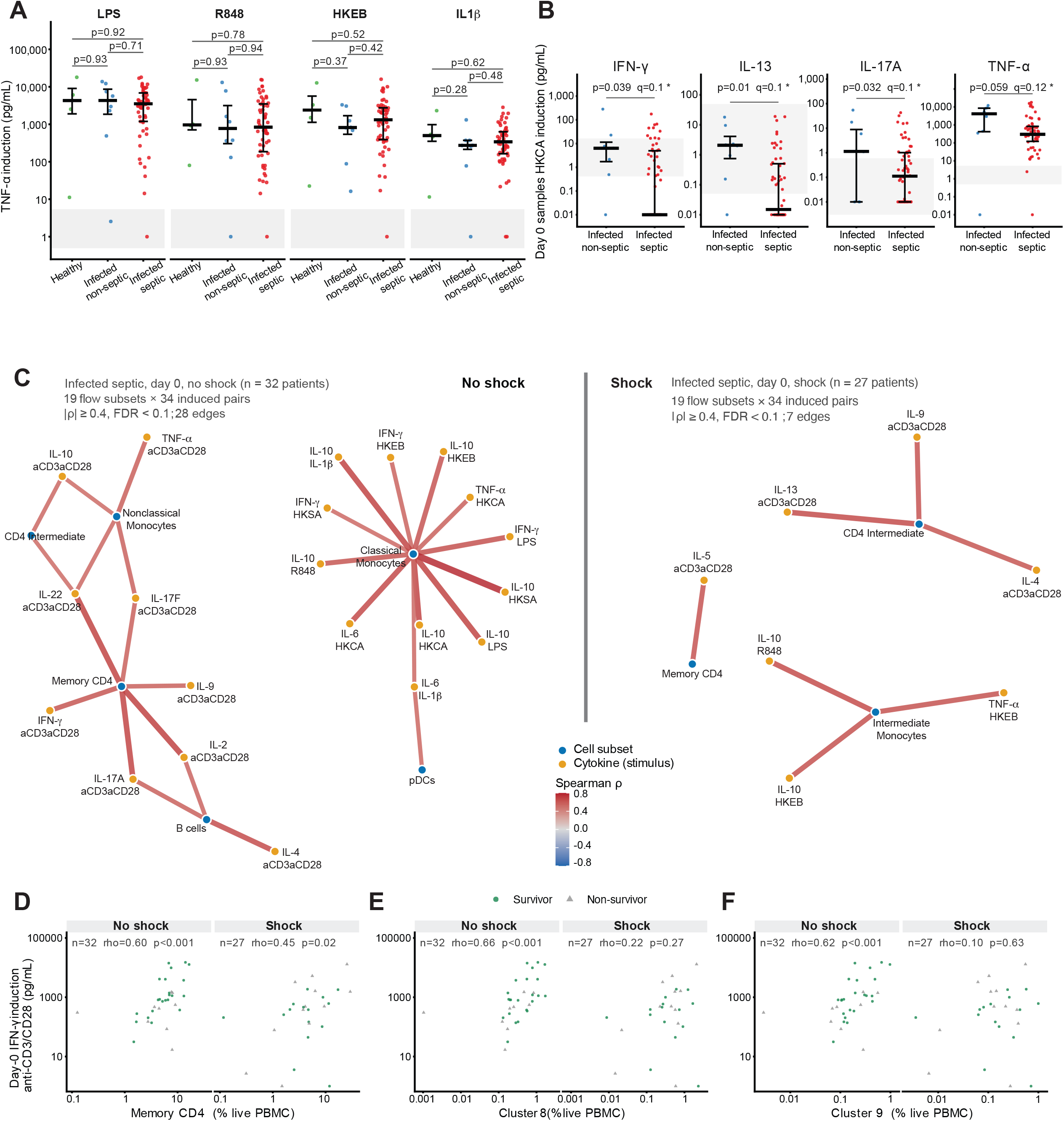
Select cytokine responses and subset-cytokine correlation network by shock status. (A) TNF-α induction by LPS, R848, HKEB, and IL-1β, stratified by healthy, infected non-septic, and infected septic groups. Unadjusted p values as indicated. (B) Select cytokine responses (IFNγ, IL-13, IL-17A, and TNF-α) to HKCA, with cytokine LLOD (lower limit of detection) range indicated by grey band. (C) Correlation network between day-0 flow subset abundances and induced cytokine responses, separately by shock stratum (19 subsets × 34 stimulus-cytokine pairs = 646 tests per stratum; edges drawn at |rho| ≥ 0.4 and FDR < 0.1). (D-F) Scatterplots of day-0 IFN-γ induction against day-0 abundance of (D) memory CD4+ T cells, (E) Cluster 8 and (F) Cluster 9, each as a percentage of live PBMCs, by shock stratum, with Spearman rho and p value as indicated. These correlations are computed across all patients in each stratum, with survival as shown by symbols (circle: survivors, triangle: non-survivors).

**Figure S6.**
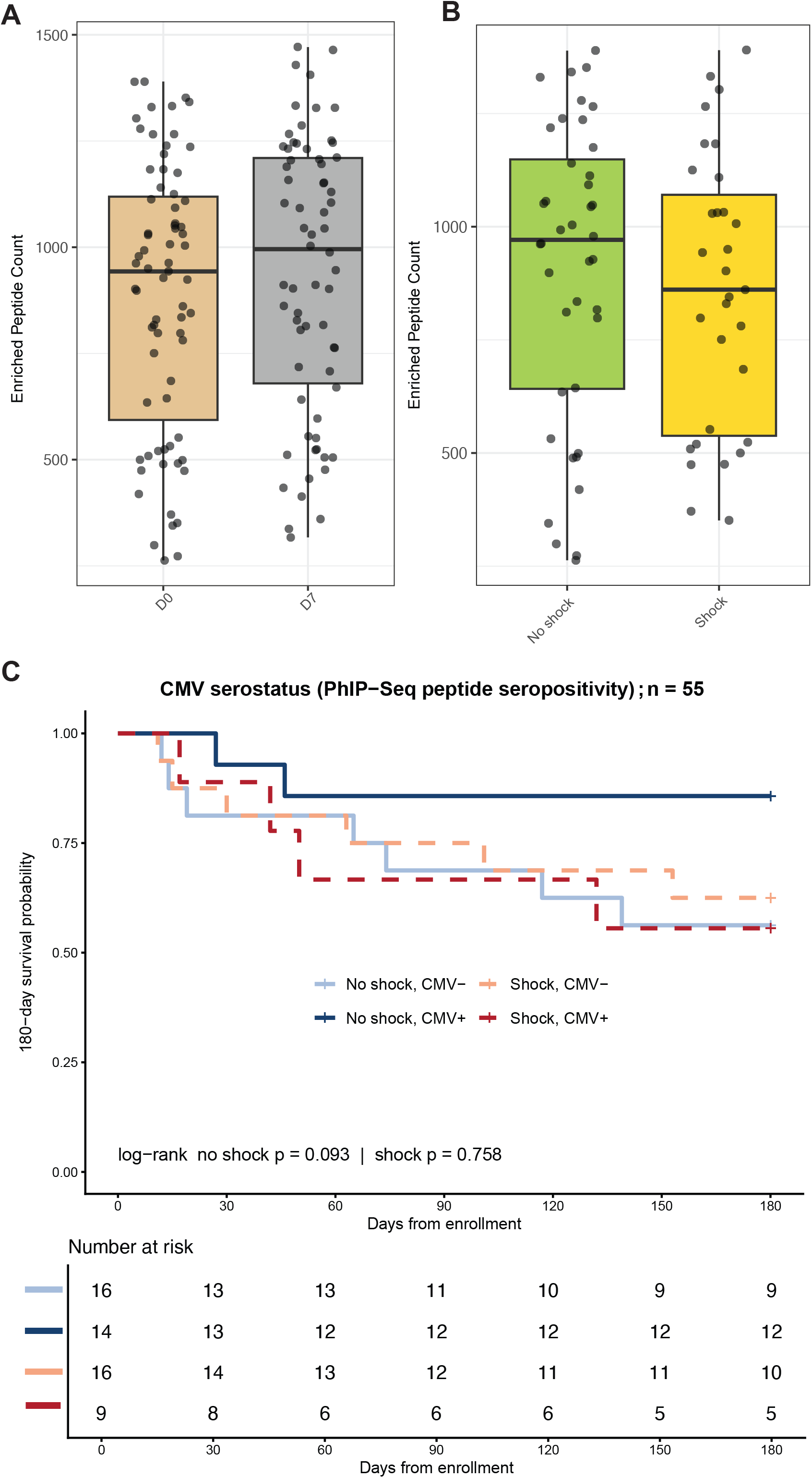
Enriched peptides by day and shock status, and CMV seropositivity Kaplan-Meier by shock status. (A) Boxplots of total enriched peptides from patients of all categories, day-0 versus day-7. Median and IQR as indicated. (B) Boxplots of total enriched peptides from infected septic patients only, no shock versus shock. Median and IQR as indicated. (C) Kaplan-Meier estimates of 180-day survival by CMV seropositivity, stratified by shock status. Sepsis without shock (solid lines), septic shock (dotted lines). The table below indicates the number of patients at risk at 30-day intervals. Statistical significance was determined by the log-rank test, p values as shown.

## Notes

### Competing Interest Statement

R.J.X. is a cofounder of Jnana Therapeutics and Convergence Bio, a scientific advisory board member at Nestle, Magnet BioMedicine and Arena BioWorks, and board director at MoonLake Immunotherapeutics; these organizations had no role in this study. The other authors declare no competing interests.

## References

1. Singer, M., Deutschman, C.S., Seymour, C.W., Shankar-Hari, M., Annane, D., Bauer, M., Bellomo, R., Bernard, G.R., Chiche, J.D., Coopersmith, C.M., et al. (2016). The Third International Consensus Definitions for Sepsis and Septic Shock (Sepsis-3). Jama 315, 801–810. 10.1001/jama.2016.0287.

2. van der Poll, T., Shankar-Hari, M., and Wiersinga, W.J. (2021). The immunology of sepsis. Immunity 54, 2450–2464. 10.1016/j.immuni.2021.10.012.

3. Giamarellos-Bourboulis, E.J., Aschenbrenner, A.C., Bauer, M., Bock, C., Calandra, T., Gat-Viks, I., Kyriazopoulou, E., Lupse, M., Monneret, G., Pickkers, P., et al. (2024). The pathophysiology of sepsis and precision-medicine-based immunotherapy. Nat Immunol 25, 19–28. 10.1038/s41590-023-01660-5.

4. Prescott, H.C., Antonelli, M., Alhazzani, W., Møller, M.H., Alshamsi, F., Azevedo, L.C.P., Belley-Cote, E., De Waele, J., Derde, L., Dionne, J.C., et al. (2026). Surviving Sepsis Campaign: International Guidelines for Management of Sepsis and Septic Shock 2026. Crit Care Med 54, 725–812. 10.1097/ccm.0000000000007075.

5. Fisher, C.J., Jr., Agosti, J.M., Opal, S.M., Lowry, S.F., Balk, R.A., Sadoff, J.C., Abraham, E., Schein, R.M., and Benjamin, E. (1996). Treatment of septic shock with the tumor necrosis factor receptor:Fc fusion protein. The Soluble TNF Receptor Sepsis Study Group. N Engl J Med 334, 1697–1702. 10.1056/nejm199606273342603.

6. Fisher, C.J., Jr., Dhainaut, J.F., Opal, S.M., Pribble, J.P., Balk, R.A., Slotman, G.J., Iberti, T.J., Rackow, E.C., Shapiro, M.J., Greenman, R.L., and, et al. (1994). Recombinant human interleukin 1 receptor antagonist in the treatment of patients with sepsis syndrome. Results from a randomized, double-blind, placebo-controlled trial. Phase III rhIL-1ra Sepsis Syndrome Study Group. Jama 271, 1836–1843.

7. Leligdowicz, A., Harhay, M.O., and Calfee, C.S. (2022). Immune Modulation in Sepsis, ARDS, and Covid-19 - The Road Traveled and the Road Ahead. NEJM Evid 1, EVIDra2200118. 10.1056/EVIDra2200118.

8. Giamarellos-Bourboulis, E.J., Kotsaki, A., Kotsamidi, I., Efthymiou, A., Koutsoukou, V., Ehler, J., Paridou, A., Frantzeskaki, F., Müller, M.C.A., Pickkers, P., et al. (2026). Precision Immunotherapy to Improve Sepsis Outcomes: The ImmunoSep Randomized Clinical Trial. Jama 335, 775–786. 10.1001/jama.2025.24175.

9. Angus, D.C. (2026). Precision Therapy for Sepsis: The End of the Beginning? Jama 335, 761–762. 10.1001/jama.2025.24099.

10. Scicluna, B.P., van Vught, L.A., Zwinderman, A.H., Wiewel, M.A., Davenport, E.E., Burnham, K.L., Nürnberg, P., Schultz, M.J., Horn, J., Cremer, O.L., et al. (2017). Classification of patients with sepsis according to blood genomic endotype: a prospective cohort study. Lancet Respir Med 5, 816–826. 10.1016/s2213-2600(17)30294-1.

11. Antcliffe, D.B., Burnham, K.L., Al-Beidh, F., Santhakumaran, S., Brett, S.J., Hinds, C.J., Ashby, D., Knight, J.C., and Gordon, A.C. (2019). Transcriptomic Signatures in Sepsis and a Differential Response to Steroids. From the VANISH Randomized Trial. Am J Respir Crit Care Med 199, 980–986. 10.1164/rccm.201807-1419OC.

12. Ganesan, A., Moore, A.R., Zheng, H., Toh, J., Freedman, M., Magis, A.T., Heath, J.R., and Khatri, P. (2025). A conserved immune dysregulation signature is associated with infection severity, risk factors prior to infection, and treatment response. Immunity 58, 2104–2119.e2105. 10.1016/j.immuni.2025.05.020.

13. Moore, A.R., Zheng, H., Ganesan, A., Hasin-Brumshtein, Y., Maddali, M.V., Levitt, J.E., van der Poll, T., Lu, J., Bouma, H.R., Scicluna, B.P., et al. (2025). A consensus immune dysregulation framework for sepsis and critical illnesses. Nat Med 31, 4084–4096. 10.1038/s41591-025-03956-5.

14. Fish, M., Rynne, J., Timms, E., Jennings, A., Lam, C., Arbane, G., Novellas, N., Campos, S., Sanderson, B., Bosco, P., et al. (2026). Temporal analyses of immune responses in sepsis reveal asynchrony between clinical stage of illness and immune states. Immunity 59, 2580–2598.e2587. 10.1016/j.immuni.2026.08.003.

15. Duffy, D., Rouilly, V., Libri, V., Hasan, M., Beitz, B., David, M., Urrutia, A., Bisiaux, A., Labrie, S.T., Dubois, A., et al. (2014). Functional analysis via standardized whole-blood stimulation systems defines the boundaries of a healthy immune response to complex stimuli. Immunity 40, 436–450. 10.1016/j.immuni.2014.03.002.

16. Brodin, P., Jojic, V., Gao, T., Bhattacharya, S., Angel, C.J., Furman, D., Shen-Orr, S., Dekker, C.L., Swan, G.E., Butte, A.J., et al. (2015). Variation in the human immune system is largely driven by non-heritable influences. Cell 160, 37–47. 10.1016/j.cell.2014.12.020.

17. Kaczorowski, K.J., Shekhar, K., Nkulikiyimfura, D., Dekker, C.L., Maecker, H., Davis, M.M., Chakraborty, A.K., and Brodin, P. (2017). Continuous immunotypes describe human immune variation and predict diverse responses. Proc Natl Acad Sci U S A 114, E6097–e6106. 10.1073/pnas.1705065114.

18. Bonecchi, R., Bianchi, G., Bordignon, P.P., D’Ambrosio, D., Lang, R., Borsatti, A., Sozzani, S., Allavena, P., Gray, P.A., Mantovani, A., and Sinigaglia, F. (1998). Differential expression of chemokine receptors and chemotactic responsiveness of type 1 T helper cells (Th1s) and Th2s. J Exp Med 187, 129–134. 10.1084/jem.187.1.129.

19. Sallusto, F., Lenig, D., Mackay, C.R., and Lanzavecchia, A. (1998). Flexible programs of chemokine receptor expression on human polarized T helper 1 and 2 lymphocytes. J Exp Med 187, 875–883. 10.1084/jem.187.6.875.

20. Brunkhorst, F.M., Engel, C., Bloos, F., Meier-Hellmann, A., Ragaller, M., Weiler, N., Moerer, O., Gruendling, M., Oppert, M., Grond, S., et al. (2008). Intensive insulin therapy and pentastarch resuscitation in severe sepsis. N Engl J Med 358, 125–139. 10.1056/NEJMoa070716.

21. Brunkhorst, F.M., Oppert, M., Marx, G., Bloos, F., Ludewig, K., Putensen, C., Nierhaus, A., Jaschinski, U., Meier-Hellmann, A., Weyland, A., et al. (2012). Effect of empirical treatment with moxifloxacin and meropenem vs meropenem on sepsis-related organ dysfunction in patients with severe sepsis: a randomized trial. Jama 307, 2390–2399. 10.1001/jama.2012.5833.

22. Larman, H.B., Zhao, Z., Laserson, U., Li, M.Z., Ciccia, A., Gakidis, M.A., Church, G.M., Kesari, S., Leproust, E.M., Solimini, N.L., and Elledge, S.J. (2011). Autoantigen discovery with a synthetic human peptidome. Nat Biotechnol 29, 535–541. 10.1038/nbt.1856.

23. Mohan, D., Wansley, D.L., Sie, B.M., Noon, M.S., Baer, A.N., Laserson, U., and Larman, H.B. (2018). PhIP-Seq characterization of serum antibodies using oligonucleotide-encoded peptidomes. Nat Protoc 13, 1958–1978. 10.1038/s41596-018-0025-6.

24. Wang, E.Y., Dai, Y., Rosen, C.E., Schmitt, M.M., Dong, M.X., Ferré, E.M.N., Liu, F., Yang, Y., González-Hernández, J.A., Meffre, E., et al. (2022). High-throughput identification of autoantibodies that target the human exoproteome. Cell Rep Methods 2. 10.1016/j.crmeth.2022.100172.

25. Hotchkiss, R.S., Monneret, G., and Payen, D. (2013). Sepsis-induced immunosuppression: from cellular dysfunctions to immunotherapy. Nat Rev Immunol 13, 862–874. 10.1038/nri3552.

26. Felmet, K.A., Hall, M.W., Clark, R.S., Jaffe, R., and Carcillo, J.A. (2005). Prolonged lymphopenia, lymphoid depletion, and hypoprolactinemia in children with nosocomial sepsis and multiple organ failure. J Immunol 174, 3765–3772. 10.4049/jimmunol.174.6.3765.

27. Hotchkiss, R.S., Tinsley, K.W., Swanson, P.E., Schmieg, R.E., Jr., Hui, J.J., Chang, K.C., Osborne, D.F., Freeman, B.D., Cobb, J.P., Buchman, T.G., and Karl, I.E. (2001). Sepsis-induced apoptosis causes progressive profound depletion of B and CD4+ T lymphocytes in humans. J Immunol 166, 6952–6963. 10.4049/jimmunol.166.11.6952.

28. Hotchkiss, R.S., Swanson, P.E., Freeman, B.D., Tinsley, K.W., Cobb, J.P., Matuschak, G.M., Buchman, T.G., and Karl, I.E. (1999). Apoptotic cell death in patients with sepsis, shock, and multiple organ dysfunction. Crit Care Med 27, 1230–1251. 10.1097/00003246-199907000-00002.

29. Schroder, K., Hertzog, P.J., Ravasi, T., and Hume, D.A. (2004). Interferon-gamma: an overview of signals, mechanisms and functions. J Leukoc Biol 75, 163–189. 10.1189/jlb.0603252.

30. Delano, M.J., Scumpia, P.O., Weinstein, J.S., Coco, D., Nagaraj, S., Kelly-Scumpia, K.M., O’Malley, K.A., Wynn, J.L., Antonenko, S., Al-Quran, S.Z., et al. (2007). MyD88-dependent expansion of an immature GR-1(+)CD11b(+) population induces T cell suppression and Th2 polarization in sepsis. J Exp Med 204, 1463–1474. 10.1084/jem.20062602.

31. Xue, M., Xie, J., Liu, L., Huang, Y., Guo, F., Xu, J., Yang, Y., and Qiu, H. (2019). Early and dynamic alterations of Th2/Th1 in previously immunocompetent patients with community-acquired severe sepsis: a prospective observational study. J Transl Med 17, 57. 10.1186/s12967-019-1811-9.

32. Stier, M.T., Sewell, A.E., Mwizerwa, E.L., Sim, C.Y., Tanner, S.M., Nichols, C.M., Durai, H.H., Jennings, E.Q., Lindau, P., Wilfong, E.M., et al. (2026). Metabolic adaptations rewire CD4(+) T cells in a subset-specific manner in human critical illness with and without sepsis. Nat Immunol 27, 236–249. 10.1038/s41590-025-02390-6.

33. Brand, A., Singer, K., Koehl, G.E., Kolitzus, M., Schoenhammer, G., Thiel, A., Matos, C., Bruss, C., Klobuch, S., Peter, K., et al. (2016). LDHA-Associated Lactic Acid Production Blunts Tumor Immunosurveillance by T and NK Cells. Cell Metab 24, 657–671. 10.1016/j.cmet.2016.08.011.

34. Milotay, G., Little, M., Watson, R.A., Muldoon, D., MacKay, S., Kurioka, A., Tong, O., Taylor, C.A., Nassiri, I., Webb, L.M., et al. (2025). CMV serostatus is associated with improved survival and delayed toxicity onset following anti-PD-1 checkpoint blockade. Nat Med 31, 2350–2364. 10.1038/s41591-025-03647-1.

35. de Brabander, J., Ong, D.S.Y., Cremer, O.L., Bos, L.D.J., Peters-Sengers, H., and van der Poll, T. (2025). Cytomegalovirus reactivation and the host immune response in patients with sepsis. J Infect 91, 106657. 10.1016/j.jinf.2025.106657.

36. Kalil, A.C., and Florescu, D.F. (2011). Is cytomegalovirus reactivation increasing the mortality of patients with severe sepsis? Crit Care 15, 138. 10.1186/cc10093.

37. Heininger, A., Haeberle, H., Fischer, I., Beck, R., Riessen, R., Rohde, F., Meisner, C., Jahn, G., Koenigsrainer, A., Unertl, K., and Hamprecht, K. (2011). Cytomegalovirus reactivation and associated outcome of critically ill patients with severe sepsis. Crit Care 15, R77. 10.1186/cc10069.

38. Unterberg, M., Ehrentraut, S.F., Bracht, T., Wolf, A., Haberl, H., von Busch, A., Rump, K., Ziehe, D., Bazzi, M., Thon, P., et al. (2023). Human cytomegalovirus seropositivity is associated with reduced patient survival during sepsis. Crit Care 27, 417. 10.1186/s13054-023-04713-1.

39. Sonnert, N.D., Rosen, C.E., Ghazi, A.R., Franzosa, E.A., Duncan-Lowey, B., González-Hernández, J.A., Huck, J.D., Yang, Y., Dai, Y., Rice, T.A., et al. (2024). A host-microbiota interactome reveals extensive transkingdom connectivity. Nature 628, 171–179. 10.1038/s41586-024-07162-0.

40. Vogl, T., Klompus, S., Leviatan, S., Kalka, I.N., Weinberger, A., Wijmenga, C., Fu, J., Zhernakova, A., Weersma, R.K., and Segal, E. (2021). Population-wide diversity and stability of serum antibody epitope repertoires against human microbiota. Nat Med 27, 1442–1450. 10.1038/s41591-021-01409-3.

41. Shrock, E.L., Shrock, C.L., and Elledge, S.J. (2022). VirScan: High-throughput Profiling of Antiviral Antibody Epitopes. Bio Protoc 12. 10.21769/BioProtoc.4464.

42. Knoll, R., Helbig, E.T., Dahm, K., Bolaji, O., Hamm, F., Dietrich, O., van Uelft, M., Müller, S., Bonaguro, L., Schulte-Schrepping, J., et al. (2024). The life-saving benefit of dexamethasone in severe COVID-19 is linked to a reversal of monocyte dysregulation. Cell 187, 4318–4335.e4320. 10.1016/j.cell.2024.06.014.

43. Quintelier, K., Couckuyt, A., Emmaneel, A., Aerts, J., Saeys, Y., and Van Gassen, S. (2021). Analyzing high-dimensional cytometry data using FlowSOM. Nat Protoc 16, 3775–3801. 10.1038/s41596-021-00550-0.

